# Allosteric Biosensors Unravel GTPase-Effector Feedback

**DOI:** 10.64898/2026.05.05.722960

**Authors:** Mingyu Choi, Roshan Ravishankar, Lihua He, Yu Yan, Gabriel Kreider, Pengning Xu, Danielle Lee, Gaudenz Danuser, Klaus Hahn

## Abstract

Fluorescent biosensors that report protein conformation *in vivo* have been invaluable for understanding how the spatio-temporal dynamics of signaling controls cells. However, for GTPases these biosensors report the activated conformation using reagents that block the binding of downstream proteins, generating dominant negative effects and altering normal cell physiology. We present here a generalizable design to make GTPase biosensors (AlloRac1 and AlloCdc42), in which a circularly permuted fluorescent protein is inserted into a conserved loop allosterically connected to the effector binding site, generating activity-dependent fluorescence without blocking ligand interactions. The Rac1 biosensor showed that effector interactions led to increased Rac1 activation, indicating an auto-regulatory positive feedback made visible by the new biosensor design. This feedback regulated the kinetics and localization of Rac1 activity, including Rac1 activity gradients that controlled motility. Feedback was generated through Rac1 interaction with the effector Pak1, which led to further activation of Rac1 by the guanine exchange factor β-Pix. The new biosensor approach enables quantitative imaging of previously obscure spatio-temporal dynamics in GTPase regulation.

## Introduction

Rho-family GTPases regulate many aspects of cell biology, including cell proliferation, cytoskeletal organization, apoptosis and gene expression. In signaling circuits they act as switches, cycling between an active and an inactive state under the influence of regulatory proteins that control both their localization and activity. The localization and kinetics of GTPase activation are often critical to downstream cellular behaviors, with spatiotemporal dynamics determining outcomes as different as apoptosis or proliferation^1,2^. Because of the importance of activation kinetics and transient localization, fluorescent biosensors have proven essential for understanding the role of GTPases^3,4^.

Almost all GTPase biosensors report activation using an affinity reagent (AR) that binds selectively to the active conformation of the GTPase. The GTPase and AR bear different fluorophores that undergo FRET when the AR binds the GTPase. AR are derived from downstream effector proteins, truncated to selectively bind active GTPase but minimize other interactions. While these biosensors have provided an important foundation for current studies, they obscure an aspect of GTPase regulation now known to play an important role in spatio-temporal dynamics – positive and negative feedback that proceeds through downstream effectors to GTPase regulatory proteins^5,6^. The effector protein fragments that are used to report GTPase activation effectively block the interactions that generate feedback.

Here we introduce an alternative biosensor strategy: It exploits a GTPase surface loop that is away from known binding interfaces for either effectors or upstream regulatory proteins, yet is allosterically coupled to conformational changes of the effector binding site^7,8^. We harness the activation-linked conformational changes of the GTPase to affect the emission of a circularly permuted green fluorescent protein (cpGFP) inserted into the allosteric loop. We generate functional biosensors for Rac1 and Cdc42 that remain responsive to upstream regulators and preserve native effector interactions. While much of a Rho family GTPase’s surface is occupied by upstream or downstream proteins, this conserved loop remains exposed and offers a general strategy to make biosensors that can interact with upstream and downstream ligands.

Using the allosteric biosensor, we directly demonstrate positive feedback for the GTPase Rac1 in live cells, and show that feedback tunes the shape and kinetics of Rac1 signaling gradients at the cell edge that control morphodynamics. The magnitude and slope of the gradients are shown to depend on Rac1’s interaction with its effector Pak1. Rac1-induced release of Pak1 from an autoinhibitory complex increases the local activation of β-Pix, a guanine exchange factor (GEF) that in turn activates Rac1.

### Development of the allosteric Rac1 biosensor

To develop GTPase biosensors that preserve downstream effector interactions, we used a previously reported analytical pipeline^7,8^ to identify surface loops that had potential allosteric connections to the effector binding site, but were removed from sites of interaction with other molecules (eg upstream regulators, downstream effectors, or the nucleotide binding site) (Fig. 1a). Our previous work had shown that one of these loops could be manipulated to control protein activity^8^, so we reasoned that its conformation might also reflect Rac1 activity.

**Figure 1.**
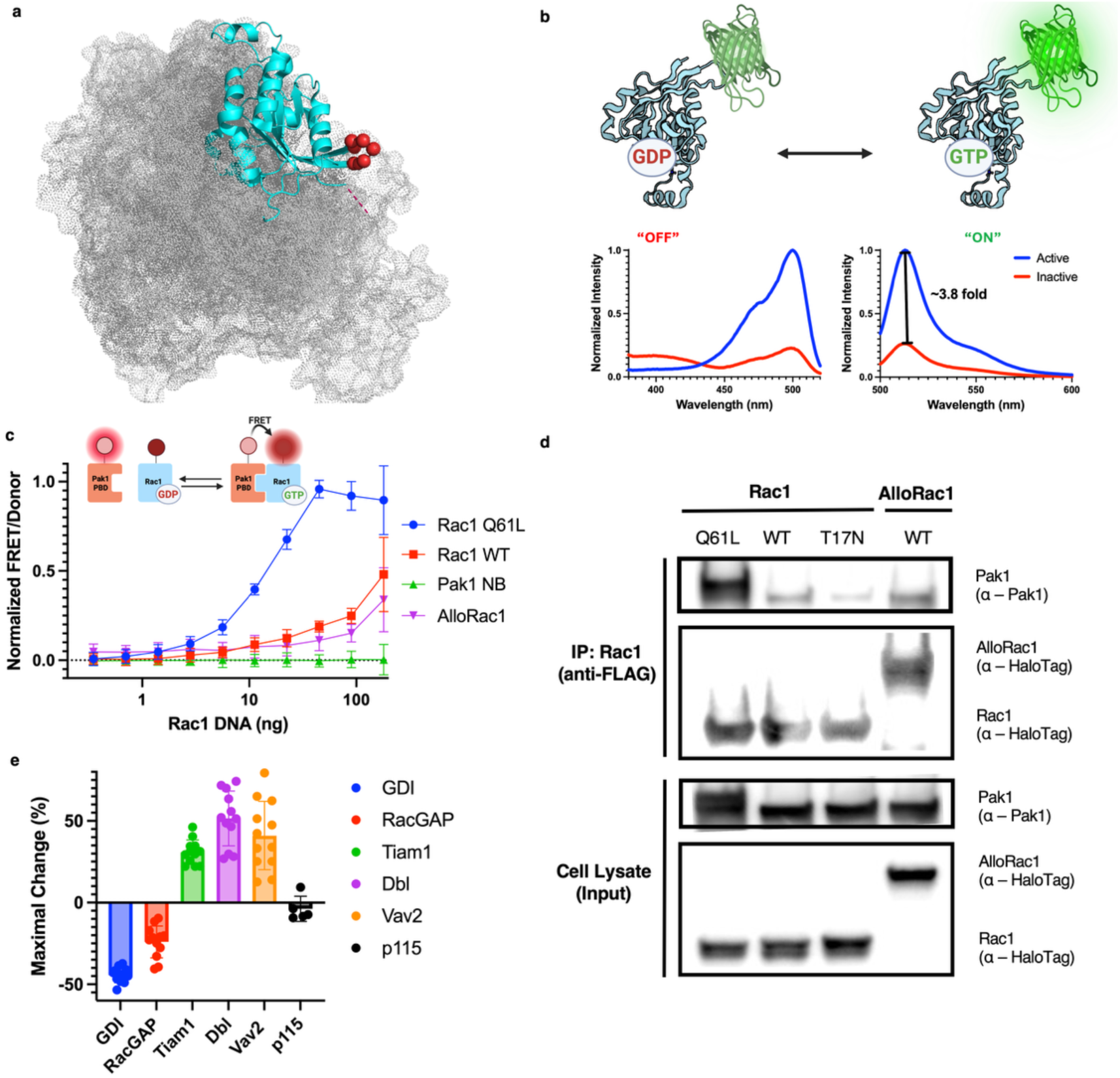
Design and characterization of the AlloRac1 biosensor. **a,** Rac1 structure (cyan) shown in complex with representative binding partners (grey; GEFs, GAPs, GDIs and effectors). The allosteric loop (red) is spatially separated from reported interaction interfaces. Structures were assembled from published co-crystal structures (PDB: 1FOE, 1HE1, 1HH4, 7AJK, 2FJU, 4YON, 1G4U, 8I5W, 6X1G, 2H7V, 7SJ4, 7USD, 2NZ8). **b,** Top, schematic of AlloRac1 design: circularly permuted GFP (cpGFP) is inserted into the allosteric loop such that its fluorescence is coupled to Rac1 activity. Bottom, excitation and emission spectra of AlloRac1 in active and inactive conformations, with peak intensity set to 1. **c,** FRET-based binding assay measuring interaction between Rac1 variants and the Pak1 p21- binding domain (Pak1-PBD). Fixed [Pak1-PBD] is titrated with varying [Rac1 levels]. (Pak1 NB = Pak nonbinding mutant, WT = wild type; N=2 independent experiments with n = 6 technical replicates per condition; error bars, s.d.) **d,** Representative co-immunoprecipitation showing association of Rac1 variants with endogenous Pak1 in HEK293T cells. **e,** Effects of upstream regulators on AlloRac1 activity. Responses are reported as maximal percent change relative to no regulator. (N = 3 independent experiments with n = 12 technical replicates per condition; for p115, N = 2 and n = 5. Error bars, s.d.)

A cpGFP whose fluorescence spectrum had responded to conformational change in other proteins^9,10^ was inserted after each residue of the allosteric loop (Fig. 1b, top; Extended Data Fig. 1). Each variant was expressed in suspended HEK293T cells, and fluorescence intensity was normalized for expression level using an N-terminal mCherry fluorescent protein on the same construct. Excitation and emission scans showed that only two insertion positions led to substantial fluorescence, one with excitation at 500 nm and the other at 405 nm. (Extended Data Fig. 1). Most insertions appeared to impair cpGFP folding or maturation.

Because the longer wavelength fluorescence was less toxic and overlapped less with cellular autofluorescence, we focused on the variant with 500 nm excitation. Linkers between cpGFP and Rac1 were optimized for dynamic range and brightness by comparing constitutively active (Q61L) and inactive (T17N) mutants. cpGFP without any added linker provided the largest activity-dependent response (Extended Data Fig. 2a). For improved live-cell imaging, we replaced mCherry with HaloTag-JF646 dye. This optimized biosensor, used for the following experiments, was named **AlloRac1**. The Q61L versus T17N mutants showed a 3.8-fold change in emission intensity (Fig. 1b, bottom) and a maximum brightness comparable to EGFP (e x QY for AlloRac1 = 31,950; EGFP=33,540^11^, Extended Data Table 1).

To assess whether AlloRac1 retained effector binding, we performed a FRET-based interaction assay. A fixed concentration of Pak1 binding domain (Pak1 residues 58–141) or its binding-deficient mutant (H83/86L; Pak1 NB, for nonbinding) was titrated with varying levels of Rac1. For FRET, the Pak1 domain was fused to mCherry and Rac1 was fused to HaloTag labeled with dye JF646. AlloRac1 behaved like wild-type Rac1, with slightly reduced FRET consistent with either weaker binding or altered positioning of the fluorophores. Constitutively active Rac1 (Q61L) produced a dose-dependent increase in FRET, whereas Pak1 NB showed no detectable response (Fig. 1c). Unlike constitutively active Rac1, wild-type Rac1 and AlloRac1 could not reach saturation within the expression range tolerated by the cells. Co-immunoprecipitation in HEK293T also showed that AlloRac1 retained Pak1 association (Fig. 1d and Extended Data Fig. 2b).

To test AlloRac1’s ability to respond to upstream regulators, we quantified its cpGFP fluorescence during titration with representative upstream molecules in HEK293T cells. Consistent with normal responses, GAP and GDI expression decreased the biosensor signal, whereas Rac1-specific GEFs increased it (Fig. 1e and Extended Data Fig. 2c). A RhoA-specific GEF (p115) produced no significant change.

Finally, to control expression levels for imaging, we generated a doxycycline-inducible PiggyBac mouse embryonic fibroblast (MEF) line expressing AlloRac1. Under low doxycycline conditions, AlloRac1 expression was substantially lower than endogenous Rac1 and did not measurably alter morphodynamic behaviors relative to the parental line (Extended Data Fig. 3a– c).

Ratiometric imaging of Q61L, WT and T17N AlloRac1 showed the expected strong differences in overall activation, and distribution of Rac1 activity consistent with previous biosensor studies (Extended Data Fig. 3d). Serum stimulation of serum starved fibroblasts increased the AlloRac1 ratio in a dose-dependent manner (Extended Data Fig. 3e,f). Together, these data establish AlloRac1 as a Rac1 biosensor that preserves effector engagement while remaining responsive to upstream regulation.

### Extension of the allosteric-loop strategy to Cdc42

To test whether the allosteric loop strategy could be applied to other Rho-family GTPases, we inserted cpGFP into the analogous loop of the GTPase Cdc42 (Extended Data Fig. 4a). Unlike Rac1, all positions produced detectable excitation at both ∼400 nm and ∼500 nm, but with greater excitation at the high energy, toxic 400 nm peak (Extended Data Fig. 4b,c). Insertion after G48 produced the greatest difference between constitutively active (Q61L) and inactive (T17N) mutants at the preferred excitation wavelength of 500 nm, so this variant was further optimized. Based on previous structure/activity studies of cpGFP^12^ we tested whether insertion of positively charged residues at the C-terminus of cpGFP could increase fluorescence at 500 nm excitation, and also improve the biosensor’s dynamic range. Adding either lysine or arginine increased the proportion of excitation at ∼500 nm (Extended Data Fig. 4d), and inclusion of a proline together with either positively charged residue enhanced this effect, with lysine-proline outperforming arginine-proline (Extended Data Fig. 4d). We referred to this optimized sensor as **AlloCdc42** (Fig. 2a). The constitutively active versus inactive mutants showed a 4.5-fold change in emission intensity (Fig. 2b, bottom) and a maximum brightness comparable to mCherry (e x QY: AlloCdc42 = 11,400, mCherry = 15,840^13^, Extended Data Table 1). Assays like those used for AlloRac1 above showed intact interactions with upstream regulators and with effectors (Extended Data Fig. 5). The presence of both 400 and 500 nm excitation bands suggested that the fluorophore was present in both a protonated and unprotonated form^14^. The added positive residues, appropriately placed through addition of proline, may favor ionization of the fluorophore to enhance 500 nm fluorescence.

**Figure 2.**
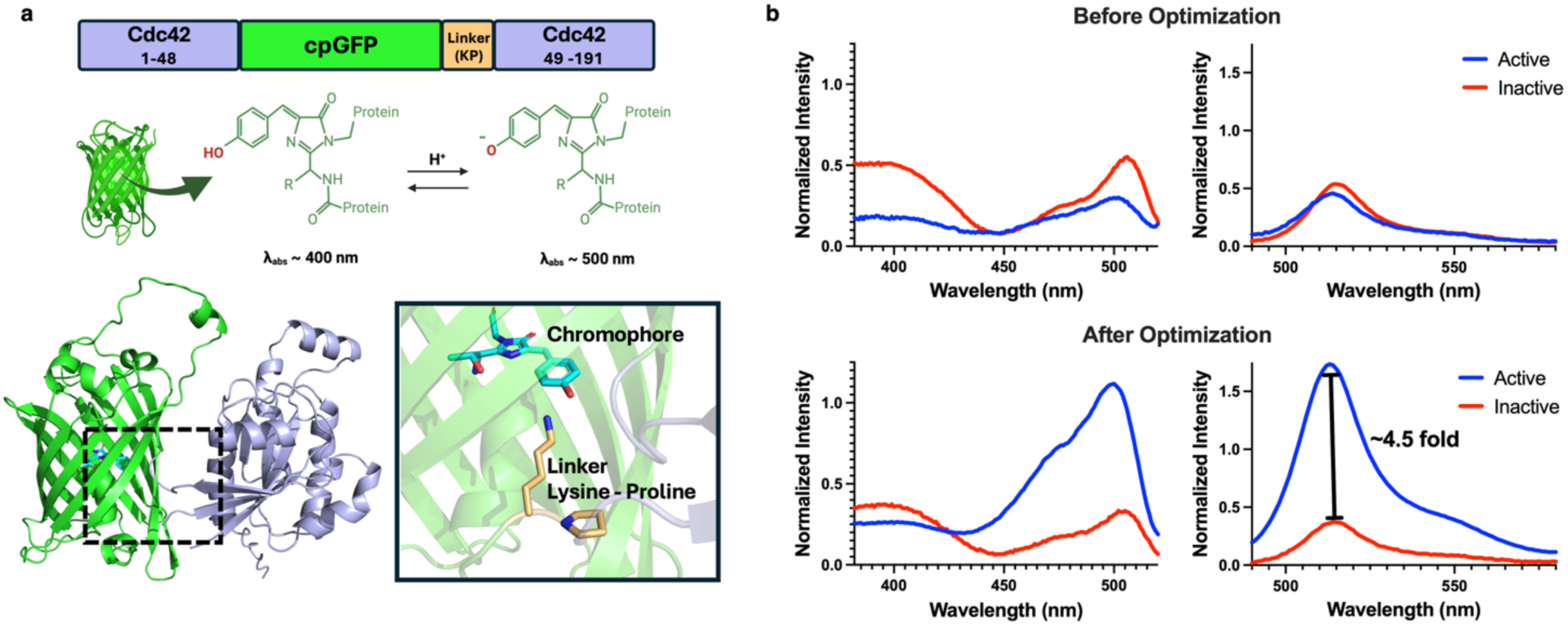
Development of the AlloCdc42 biosensor. **a,** AlloCdc42 structure and fluorophore, showing the ionizable oxygen that affects excitation peaks (top). Predicted structure of AlloCdc42 generated with AlphaFold (bottom). Inset, interface between Cdc42 (purple) and cpGFP (green). The two residues closest to the GFP fluorophore are highlighted (orange). **b,** Normalized excitation and emission spectra of AlloCdc42 in constitutively active and inactive states before (top) and after (bottom) optimization.

We generated a doxycycline-inducible PiggyBac MEF line expressing AlloCdc42 and tuned induction to keep biosensor levels below endogenous Cdc42 (Extended Data Fig. 6a,b). Q61L, WT, and T17N mutants of AlloCdc42 reported the expected level and distribution of activity, and the fluorescence ratio of WT AlloCdc42 increased upon serum stimulation (Extended Data Fig. 6d–f). Interestingly, in cells showing a clear direction of movement, Cdc42 activity was more focused at the leading edge, consistent with Cdc42’s reported role in determining the direction of migration (Extended Data Fig. 6c).

Successfully generating biosensors for both Rac1 and Cdc42 showed that allosteric loop targeting can provide a generalizable route to GTPase biosensors.

### Rac1 effector engagement controls Rac1 activity and affects protrusion dynamics

Accumulating evidence suggests that downstream signaling from Rac1 can modulate Rac1 activity, either positively or negatively^15–18^. With AlloRac1 we could test this in live cells, and examine whether such feedbacks play a role in Rac1’s control of protrusions, a dynamic cell behavior that requires localized and precisely timed Rac1 activation. We first mutated the biosensor to perturb interactions with downstream effectors (T35S, Fig. 3a)^19^. Even though the biosensor was present in only tracer amounts, mutating the biosensor was sufficient to reduce the mean Rac1 activity of the cell and to affect protrusion dynamics (Fig. 3b). The mutation caused the cell to spend a smaller percentage of its time undergoing protrusion, and also affected the proportion of time the cell spent in directed versus random migration, a property previously attributed to Rac1 signaling (Fig. 3c). In contrast, there was no apparent effect on the percent of time spent in retraction. The remarkable potency of mutating only trace amounts of GTPase suggested that the effects were either localized, or signaling from effectors was amplified downstream.

**Figure 3.**
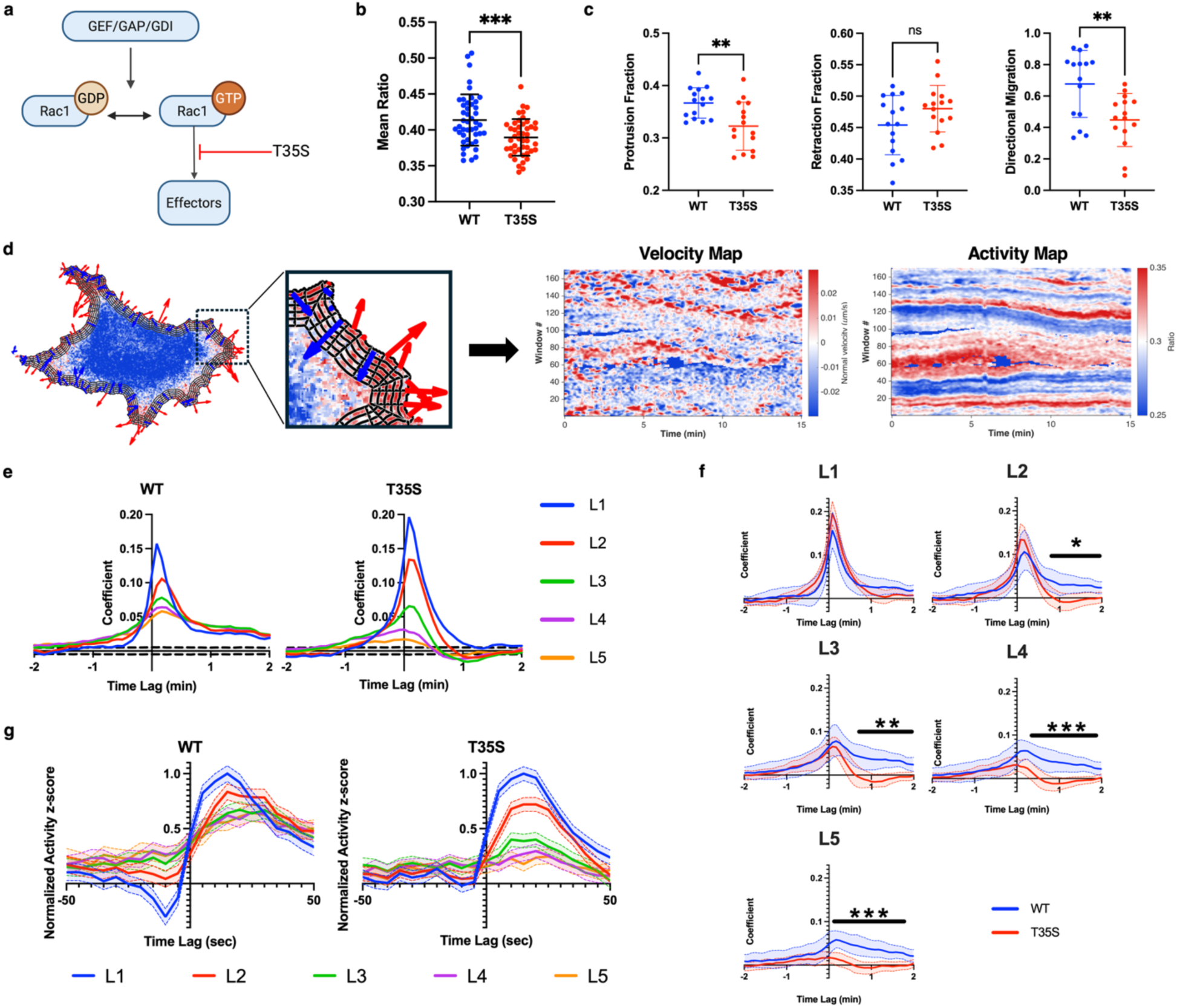
Effector engagement influences the spatiotemporal dynamics of Rac1 activation and its coupling to edge movements. **a,** Schematic showing the Rac1 GDP–GTP cycle, its control by GEFs, GAPs and GDIs, and effector binding by GTP-loaded Rac1. T35S mutation broadly disrupts effector interactions. **b,** Whole cell mean AlloRac1 ratio for WT and T35S mutant (N = 3 independent experiments with n = 45 cells per condition, lines indicate population means with error bars s.d. Two-tailed Welch’s T-test; ***P ≤ 0.001). **c,** Protrusion fraction (fraction of time spent in protrusion), retraction fraction (fraction of time spent in retraction), and directional migration (measure of directed migration = net displacement /total displacement) quantified per cell for WT and T35S. Two-tailed Welch’s t-test ns, P > 0.05; **P ≤ 0.01. **d,** Windowing approach used to correlate cell edge dynamics and the associated Rac1 activity. Rac1 activity was sampled over time in windows at the cell edge, in layers of 0.86 µm depth, and the corresponding edge velocity was tracked simultaneously. Arrows show magnitude and direction of edge velocity, colors indicate Rac1 activation level. **e,** Cross-correlation between Rac1 activity and edge velocity computed separately for each layer; curves show the population mean (N = 3 independent experiments with n = 15 cells per condition). Dotted area indicate 95% permutation-based confidence level from time-shuffled data (1000 times). **f,** Across layers, WT vs T35S differences were evaluated using a two-sided cluster-based permutation test, at correlations with lags of 0–2 min (cluster-forming threshold: pointwise *P*<0.05; minimum cluster size = 3 points; 5,000 permutations). Asterisks indicate cluster-level FWER-corrected *P* values (*P ≤ 0.05; **P ≤ 0.01; ***P ≤ 0.001). Shaded bands indicate mean ± 95% confidence level. Most significant effects were in the tail of the correlation curve, at time lags from ∼1–2 minutes in layers 3, 4 and 5. **g,** WT vs T35S Rac1 activity reported in window-based Z-score relative to protrusion onset (t = 0). Values were normalized to the maximum of the first layer. Shaded bands indicate mean ± 2x s.e.m.

### Rac1-effector interactions tune the spatiotemporal dynamics of Rac1 activity gradients

These studies indicated an effector-mediated feedback interaction controlling Rac1 activity. We sought to verify and dissect the role of this feedback in Rac1 control of morphodynamics. It is well established that Rac1 controls cell protrusion, and the timing and position of Rac1 activation at the cell edge correlates with protrusion initiation, velocity, duration and other parameters of morphodynamics^20–22^. We used AlloRac1 to examine how Rac1– effector interactions impact the kinetics and localization of Rac1 and correlated cell edge behaviors. The velocity of the cell edge and nearby Rac1 activity were simultaneously quantified in sampling windows at different distances from the edge^20,23^ (Fig. 3b). Consistent with published biosensor studies^20–22^, Rac1 activity was coordinated with edge motion most strongly at the cell boundary, and became progressively less coupled for windows deeper into the cell (Fig. 3e and Extended Data Fig. 7a). Coupling was maximal ∼5–10 s after a change in edge velocity, consistent with Rac1 activity reinforcing protrusion^20,24^. Rac1 activity remained correlated with edge movements after they occurred, producing a sustained tail in the cross-correlation curves. Disrupting effector engagement with T35S reduced this sustained correlation while largely preserving the timing and localization of the peak occurring immediately at the cell edge (Fig. 3e and Extended Data Fig. 7a). The loss of sustained correlation was most pronounced ∼2–4 µm from the cell edge. A cluster-based permutation test (Fig. 3f) and comparison of the areas under the correlation curves confirmed that the greatest effects on edge–Rac1 correlation occurred for windows deeper into the cell (Extended Data Fig. 7b). Notably, the T35S cross-correlation curves closely resembled those from previously published biosensors where the effector binding site was blocked by an affinity reagent, preventing interaction with endogenous effectors^21,22^.

These results suggested that effector engagement is dispensable for the initial edge-proximal Rac1 response but contributes to maintaining Rac1 activity over longer timescales and at positions farther from the cell edge, potentially through positive feedback. To test this idea, we quantified Rac1 activity in individual protrusions before and after protrusion onset (Fig. 3g). Consistent with the cross-correlation analysis, the deeper into the cell we sampled, the greater the reduction in Rac1 activation caused by T35S.

### PAK and β-Pix mediate the Rac1 effector feedback loop

We sought to determine which molecules were mediating the effector-driven feedback loop. To test this, we first examined Rac1 point mutations known to either broadly disrupt interactions with downstream proteins (T35S), or to more selectively reduce binding to effectors that contain the CRIB domain (Y40C; CRIB = Cdc42/Rac–interactive binding; Fig. 4a). Rac1 activation was reduced by T35S, as expected from the results above, but was even more strongly reduced by Y40C (Fig. 4d), suggesting that a CRIB-domain effector plays an important role in feedback. The weaker effect of T35S may reflect its broader disruption of multiple downstream pathways, including both positive and negative circuits, whereas Y40C may more selectively impair a positive feedback mechanism that is CRIB-dependent. We chose to focus on Pak1, a canonical CRIB-domain effector implicated in regulation of Rac1 signaling^25^. We used two small molecule inhibitors that disrupt Pak signaling in different ways (Fig. 4c). In the absence of Rac1, Pak1 forms an autoinhibitory dimer in which Pak1 kinase activity is suppressed, while Pak1 remains associated with partner proteins such as the Rac1 GEF b-Pix (Fig. 4b). The inhibitor IPA-3 binds to the dimer, preventing Rac1-mediated dissociation, thereby blocking the release of Pak1 and the associated GEFs. NVS-Pak1, in contrast, allosterically inhibits Pak1 kinase activity. IPA-3 significantly reduced overall Rac1 activity, while NVS-Pak1 had no effect (Fig. 4d), indicating that PAK was important to the feedback loop, but that its kinase activity was not strongly involved.

**Figure 4.**
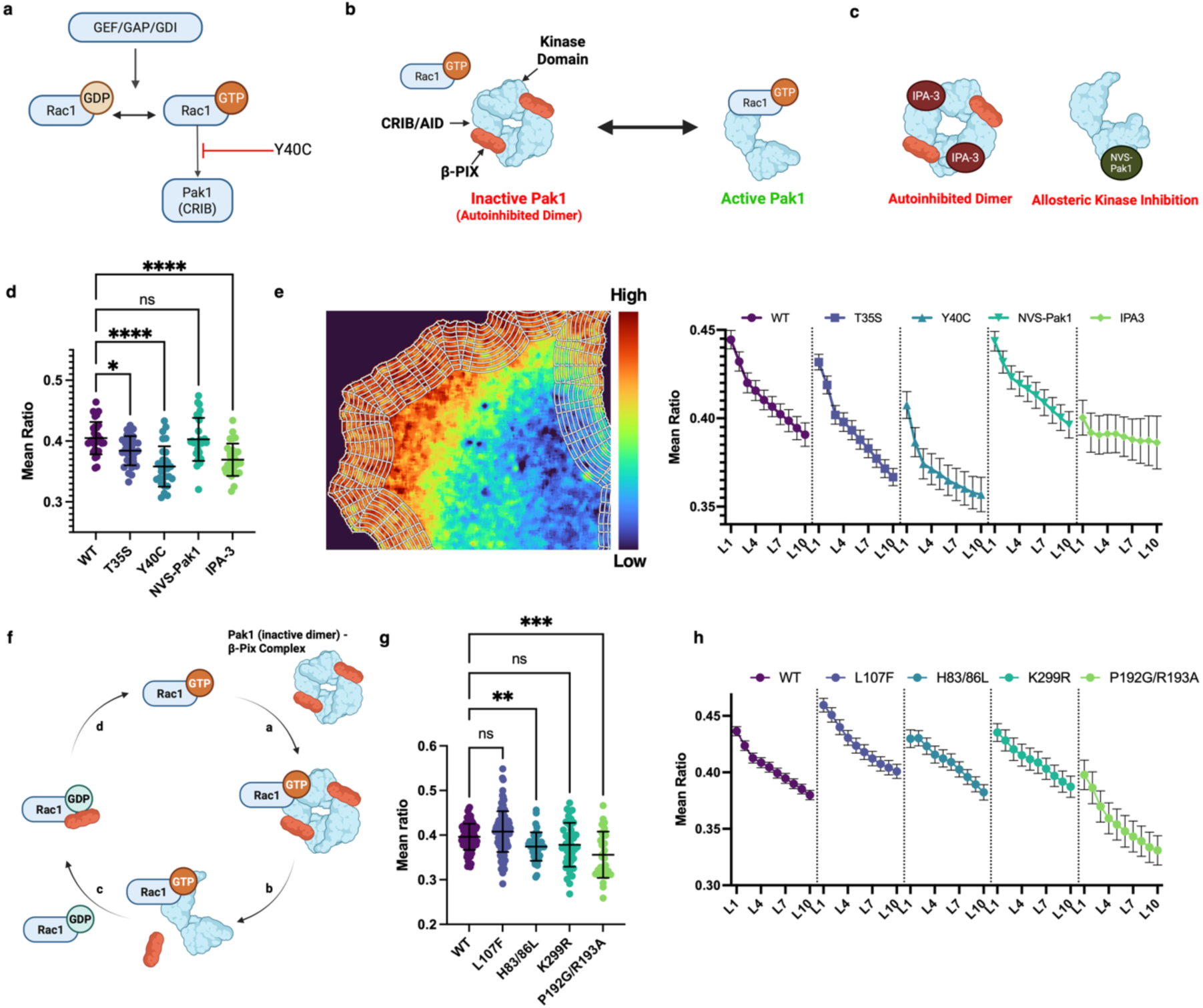
Rac1-Pak1 engagement modulates the Rac1 activity gradient through β-Pix. **a,** Schematic of the Rac1 GDP–GTP cycle. Y40C mutation disrupts interactions with effectors containing the CRIB-domain, including Pak1. **b,** Model of Pak1 regulation. Pak1 is autoinhibited when in a homodimer complex also containing β-Pix. Pak1 binding to Rac1–GTP relieves autoinhibition and releases binding partners, including β-Pix. **c,** Pak1 inhibitors used in this study: NVS-Pak1 allosterically inhibits kinase activity, IPA-3 stabilizes the inactive, autoinhibited Pak1 state. **d,** Rac1 activity quantified as the whole cell mean biosensor ratio, for Rac1 mutations and Pak1 inhibitor treatments. Data represent (N = 3 independent experiments with n = 30 cells per condition; Lines indicates population mean with error bars s.d. Ordinary one-way ANOVA Dunnett’s multiple comparisons test; ns, P > 0.05; *P ≤ 0.05; ****P ≤ 0.0001). **e,** Sampling windows for profiling Rac1 activity relative to the cell edge. Layer 1 (L1) is nearest the edge and successive layers extend inward. Each layer spans ∼*0.86 μm*. (Points show the population mean ±s.e.m.; same population as in d). **f,** Schematic of the Rac1–Pak1 positive-feedback loop. (a): Rac1–GTP binds Pak1 (b): this promotes dissociation of Pak1 and release of the Rac1 GEF β-Pix. (c): Local Rac1-GDP binds GEF β-Pix (d) this in turn activates additional Rac1 to sustain Pak1 activation. **g,** Whole cell mean Rac1 activity for cells expressing Pak1 mutants meeting the selection criteria (Data collected over 10 days. n = 105 (WT), 104 (L107F), 42 (H83/86L), 44 (K299R), 28 (P192G/R193A). Line indicates population mean with error bars s.d. Kruskal-Wallis test; ns, P > 0.05; **P ≤ 0.01; ***P ≤ 0.001.) **h,** Profiles of Rac1 activity gradients, determined as in (**e**). Points show the population mean ± s.e.m. (same population as in g).

We asked how these mutations and inhibitors affected the Rac1 activity gradient at the cell edge (Fig. 4 e). Both Y40C and IPA-3 reduced the magnitude of the activity gradient, and IPA-3 also flattened the gradient. Again, NVS-Pak1 had minimal effect. Together, these results strongly suggest that Pak1 contributes to the activating feedback not through its kinase activity, but by acting as a scaffold.

We reasoned that Rac1 activation might result from release of β-Pix when Rac1 disrupts the Pak1 autoinhibitory complex (Fig. 4f)^18,26,27^. To test this, we expressed the following Pak1 mutants in AlloRac1 cells: L107F, which produces a constitutively active Pak1^28,29^; H83,86L, which blocks Pak1-Rac1 binding^30^; K299R, which abrogates Pak1 kinase activity^31^; and P192G/R193A, which reduces β-Pix binding to Pak1^26^. Control experiments established an expression range where exogenous Pak1 did not by itself alter overall Rac1 activity (Extended Data Fig. 8). Consistent with our hypothesis, the mutation that reduced Pak1–β-Pix binding produced the strongest reduction in Rac1 activity, supporting the idea that β-Pix release from the autoinhibitory complex contributes to Rac1 activation. Abrogating the ability of Rac1 to bind to Pak1 also significantly reduced activation. Importantly, inhibiting or increasing kinase activity had no significant effect (Fig. 4g). Similar to these effects on overall Rac1 activity mutations that disrupted the Pak1– β-Pix interaction had the strongest effect on the cell edge gradient, whereas the other mutants had minimal effects (Fig. 4h).Together, these results support a model in which Rac1 creates positive feedback via disruption of the Pak1 autoinhibitory complex, with release of β-Pix, and not through Pak1 kinase activity.

Finally, we asked whether Rac1 signaling through Pak1 and β-Pix affected the correlation between Rac1 activity and protrusion events (Extended Data Figs. 9 and 10). The cluster-based permutation test used in Fig. 3f above showed that IPA-3, which inhibits Rac1’s activation of Pak1 (Fig. 4a), did affect the correlation of Rac1 activity and edge dynamics, but more weakly than did the Rac1 T35S mutation, which broadly blocks effector interactions. This contrasts with effects on gradient shape and magnitude (Fig. 3f above), where IPA-3 had a stronger effect than T35S. Both T35S and IPA-3 exerted their effects on the tail of the correlation curve, at longer times after edge behaviors. No other mutants or inhibitors had a significant effect. Together, these data show that Rac1-Pak1 interaction strongly affects the spatial distribution of Rac1 activity, but more weakly affects the kinetics of activation relative to edge dynamics, and that the kinase activity of Pak1 is not critical to either. Control of kinetics appears to involve multiple effectors.

## Discussion

We described here a design for small GTPase biosensors that harnesses allosteric coupling between an exposed surface loop and the effector binding site. A fluorescent protein inserted in the loop responded to effector site conformational changes but did not sterically perturb interactions with upstream regulators or downstream effector proteins. This enabled the biosensor to reveal Rac1 control of its own spatio-temporal dynamics, via effector feedback pathways. A positive feedback loop was important in shaping the gradient of Rac1 activity that controls cell edge morphodynamics. The overall shape of this gradient was found to be modulated specifically by Rac1 unlocking the Pak1 autoinhibitory dimer and releasing the Rac1 activator β-Pix.. In contrast, the timing of cell edge Rac activation was not so predominantly controlled by this one effector.

Our success in building allosteric biosensors for both Rac1 and Cdc42 bodes well for broad applicability. The surface loop where cpGFP was inserted is found throughout the small GTPases, with a well preserved connection to the effector binding site via two strands of a beta pleated sheet. The new design does not produce dominant negative effects caused by effector site occupancy, and engineering biosensors for other GTPases will be simpler because a single, responsive fluorescent protein is used rather than a FRET pair. Limitations remain--AlloSensors bind effectors with reduced affinity relative to wild-type, but we could tune the tradeoff between dynamic range and effector Kd using linker mutations that positioned positive residues near the cpGFP fluorophore.

Mutation of only the small, tracer amount of biosensor in our experiments was enough to strongly affect signaling and morphodynamics. Intriguingly, biosensor mutations that disrupted interactions with downstream effectors significantly reduced biosensor activation, showing that effector-mediated feedback signals impacted the actual Rac molecules that had bound the effectors. The precise mechanism for such highly localized feedback amplification of Rac-signaling remains to be determined, but our data suggests that signals remain local because of a clustering of Rac1, Pak1 and the Rac1 activator β-Pix in complexes, via a scaffold and/or membrane curvature ^32,33^. It is also possible that downstream amplification produces a large number of molecules capable of diffusing back and impacting the biosensor.

In future work, allosteric GTPase activity reporters can be combined with fluorescent analogs of effectors to examine the localization and kinetics of specific signal-effector interactions. This can be combined with optogenetic control of upstream molecules^8,34–36^. The biosensor design presented here can be a stepping stone towards dissection of dynamic effector-driven feedback architectures and their role in living cells.

## Methods

### AlloSensor Design

Allosteric loops were identified for GTPases using the previously described computational pipeline^7,8^. In brief, surface exposed loops were identified using crystal structures and computationally calculating solvent accessible area. To minimize functional perturbation, loops with highly conserved sequences were eliminated from consideration. Contact maps were generated using crystal structures and homology models. Loops showing a continuous series of close proximity residues stretching to the effector binding site were tested for allosteric coupling. Circularly permuted GFP (Addgene#40753)^10^ was inserted after each amino acid in the allosteric loop using Gibson Assembly Master Mix (New England Biolabs, NEB). Fluorescence excitation/emission spectra were used to identify promising insertion sites. Constitutively active (Q61L) and inactive (T17N) mutants were used to gauge the dynamic range of the variants. For linker optimization, mutants were made using Q5 Hot Start High-Fidelity Master Mix (NEB), DpnI (NEB), T4 PNK (NEB), and T4 DNA Ligase (NEB).

### Spectral analysis of AlloSensors

HEK293T cells were plated on 12-well plates and cultured to 60-70% confluency. Cells were transfected with biosensors using Lipofectamine (Invitrogen) and Plus reagent (Invitrogen) using the manufacturer’s protocol. On the next day, growth medium supplemented with 200 nM JF646 dye (Janelia) was added and cells were incubated for 2 hours. Cells were suspended in cold PBS (Corning) supplemented with 1% Fetal Bovine Serum (FBS, Gemini) and 10 mM HEPES buffer (Corning). The cells were spun down and resuspended in the same medium to wash away excess dye. Corrected fluorescence spectra were obtained using a Fluorolog fluorometer (Horiba). For GFP, emission was fixed at 530 nm and excitation was collected from 350-520 nm. For the emission spectra, samples were excited at 480 nm and emission was collected from 500-560 nm. Spectra were normalized for expression level using a second fluorophore attached to each protein as described in the results section. “Brightness” was defined as the product of extinction coefficient and quantum yield. For the biosensors, brightness was determined by comparing biosensor fluorescence to a reference fluorophore with known brightness expressed in the same cell, normalized for the use of different excitation and emission wavelengths.

### High-content microscopy screening

Biosensor interactions with upstream regulators and downstream effectors were probed via high-content imaging as previously described^37^. Briefly, HEK293T cells were plated in 96-well plates with flat glass bottoms (Greiner) coated with poly-L-lysine (Sigma). Cells were transfected using Lipofectamine (Invitrogen) and Plus reagent (Invitrogen) with biosensor DNA together with a library of regulators or downstream effectors. Regulator or effector DNA was titrated against a constant amount of biosensor DNA in a separate 96-well, u-clear plastic bottoms (Greiner) to assemble transfection complexes, which were then applied to the cells. The next day, cells were incubated for 2 hours in growth medium supplemented with 200 nM JF646 dye (Janelia). Medium was replaced with PBS containing 1% FBS and 10 mM HEPES (Corning) immediately before imaging. Images were acquired on an Olympus IX-81 inverted microscope equipped with a 10X/0.4 NA Olympus objective, mercury arc illumination, and MetaMorph screen acquisition software (Molecular Devices). JF646 images were collected using filters with excitation FF01-640/14 and emission ET705/72; GFP images were collected using excitation HQ470/40X and emission FF01-530/43 with an FF405/496/593/649-Di01 dichroic mirror. Images were recorded on an Orca Flash4 sCMOS camera (Hamamatsu) and analyzed using MATLAB (MathWorks). For each well, four fields of view were acquired and the summed intensity was computed for each channel. Background was estimated from mock-transfected wells and subtracted from each measurement. For titration experiments with upstream regulators, per-well biosensor signal was quantified as the ratio of summed GFP intensity to summed JF646 intensity and normalized to the cells expressing biosensor alone (no regulator). Regulators tested were RhoGDI (full length), Dbl (495–826), Vav2 (191–573), Tiam1 (C1199), RacGAP1 (full length), p115RhoGEF (full length), Asef (full length), and Intersectin (1229–1586). For the FRET binding assay, bleedthrough coefficients were estimated from wells expressing a nonbinding Pak1 mutant and used to correct measured FRET prior to reporting the FRET ratio.

### Co-Immunoprecipitation

HEK293T cells were seeded on 6-well plates and transfected at 70% confluency with AlloSensor constructs using Lipofectamine and Plus reagent (Invitrogen). At 18 hr post-transfection, growth medium was aspirated and the cells were washed with 2 mL of PBS twice. To each well, 300 uL of IP lysis buffer (25 mM Tris-HCL pH 7.4, 150 mM NaCl, 1 mM EDTA, 1% Igepal CA-630, and 5% glycerol) containing protease inhibitor (cOmplete EDTA-free, Roche) and phosphatase inhibitor (PhosSTOP, Roche) were added. The lysate was transferred to pre-chilled Eppendorf tubes on ice. The tubes were rotated for 30 minutes in the cold room for complete lysis. Following incubation, the lysate was centrifuged at 10,000 x g for 10 minutes at 4°C. After centrifugation, 30 uL of supernatant was removed from each sample to assess protein expression. The supernatant was mixed with 4X Laemmli sample buffer (Bio-Rad) with 10% 2-mercaptoethanol (Sigma), boiled for 5 mins and stored at −20°C for Western blotting. Anti-FLAG M2 agarose beads (Sigma) were blocked with 1% BSA in IP buffer overnight at 4°C. The beads were then washed three times with IP lysis buffer. To each sample, 40 uL of anti-FLAG M2 agarose beads were added and placed on a rotating mixer at 4°C for 2 hrs. Following incubation with anti-FLAG agarose beads, the samples were washed three times with IP lysis buffer. To elute the sample, 20 uL of 0.2 mg/mL 3x FLAG peptide (APExBio) solution in TBS was added to each sample and placed on a rotating mixer at 4°C for 25 mins. The eluant was collected by piercing a hole in the cap and the bottom of the tube with an 18-gauge needle, placing the tube in another labeled Eppendorf tube, and then centrifuging at 1000 x g for 1 min. 10 uL of 4x Laemmli sample buffer (10% 2-mercaptoethanol) was added. The sample was then boiled for 5 mins and stored at −20°C.

### Western Blotting

Equal amounts of protein were loaded into the wells of 5-15% gradient SDS-PAGE gel (Bio-Rad) with molecular weight markers. The gel was run at 120 V until the bands were fully separated. PVDF membrane was activated in methanol and washed with transfer buffer prior to preparing a transfer stack. Protein was transferred to the membrane using the Trans-Blot Turbo Transfer System (Bio-Rad). The membrane was blocked in 5% non-fat dry milk in TBS-T for an hour on a shaker at room temperature. The membrane was then washed three times with TBS-T buffer with 10 minute incubation each time. Primary antibody (see antibody list below) in 3% BSA in TBS-T was applied to the membrane and incubated overnight on a shaker at 4°C. The following day, the membrane was washed three times with TBS-T buffer with 10 minute incubation each time. Secondary antibody (Jackson Laboratories, see reagent section) was diluted in 5% non-fat dry milk in TBS-T (1:10,000), then applied to the membrane and incubated for an hour at room temperature on a shaker. The membrane was washed again as described above before being imaged with Clarity ECL substrate (Bio-Rad) on the imager.

### Antibodies Used

HaloTag (Promega, G9281), Pak1 (Cell Signaling, 2602S), Rac1 (Cell Signaling, 2465), FLAG M2 (Sigma, F1804-5MG), Cdc42 (Cell Signaling, 2562S), anti-Rabbit IgG (Jackson ImmunoResearch, 211-032-171).

### Stable Cell Line Generation

Stably-expressing tet-off MEF cells (Clontech) were seeded in six-well plates and cultured in growth medium until reaching 70–80% confluency. Piggybac vectors containing AlloSensors were co-transfected with transposase using a 4:1 ratio (Piggybac DNA:Transposase DNA) and TransIT-X2 (Mirus Bio) following the manufacturer’s protocol. Eighteen hours post-transfection, the medium was replaced with fresh growth medium supplemented with 1 µg/mL doxycycline and antibiotic. The antibiotic concentration was doubled every two days until a maximum concentration was reached. For puromycin selection, starting concentration was 1 µg/mL with a maximum concentration of 10 µg/mL. For hygromycin, starting concentration was 50 µg/mL with a maximum concentration of 500 µg/mL. Cells were passaged and maintained under selection for approximately 2 weeks. In the AlloRac1 MEF cell line, this yielded expression in more than 90% of cells. For other cell lines, fluorescence-activated cell sorting (FACS) was used to ensure expression. Cells were aliquoted and frozen in growth medium supplemented with 7.5% DMSO and 1 µg/mL doxycycline. Freezing was performed at –80 °C for one week before transferring aliquots to liquid nitrogen for long-term storage. Because we were concerned that expression might decline over time, cells beyond passage 15 were maintained in antibiotic supplemented media or re-sorted by FACS.

### Live Cell-Imaging

Stable cell lines were maintained in complete growth medium supplemented with 20 ng/mL doxycycline. To induce expression, cells were washed three times with growth medium, with a minimum of 5 min incubation between washes. After washing, growth medium with 20 pg/mL doxycycline was added to the cells. The day before imaging, cells were plated on coverslips coated with 5 µg/mL of fibronectin (Sigma). On the day of imaging, cells were incubated with growth medium containing 200 nM JF646 for two hours. Before imaging, media was replaced with FluoroBrite (Gibco) containing 7.5% FBS, 1X GlutaMax (Gibco), 20 mM HEPES (Corning), and 20 pg/mL doxycycline.

For Pak1 inhibition experiments, the imaging medium was supplemented with either 1 µM NVS-Pak1 (Wolff Lab) or 5 µM IPA-3 (MedChem) and cells were incubated with inhibitor for 2 hours prior to imaging.

Images were acquired on an Olympus IX-81 inverted microscope with an open heated chamber (Warner Instruments) using either a UPlanAPO 40x Silicone 1.25 NA or UPlanAPO 60x Oil 1.5 NA objective. MetaMorph software was used to control the microscope and acquire images. For illumination, a 100 Watt mercury arc lamp was used with either 1% or 3% ND filter for both JF646 and GFP channels. Filters and dichroic mirror were identical to the ones described in the previous section (High-content microscopy screening). For time series data, images were acquired every 5 seconds for 15 min.

### Image processing and analysis

GTPase activity levels were measured in living cells by adapting a previously reported analysis pipeline (https://github.com/DanuserLab/u-probe)^38^. Briefly, we monitored the ratio of GFP intensity to the normalizing fluorophore, HaloTag-JF646, on a pixel by pixel basis. Camera dark current noise was determined and subtracted by obtaining images without excitation. To correct for uneven illumination, we took images of a uniform dye solution using the identical imaging condition for each wavelength at multiple stage positions. These images were averaged and then normalized to create a reference image with an average intensity of one. Dark current corrected images were divided by this shade reference image. Shade corrected images were cropped and then segmented using ‘MSA Segmentation’^39^. To measure background fluorescence at each frame, the cell mask was dilated and the original cell mask was then subtracted from the dilated mask to create a background cell mask. The intensity of this region was subtracted from each channel at every frame. To create GTPase activity images, the background-subtracted GFP channel was divided by the background subtracted HaloTag-JF646 channel. The cell mask was then applied to this ratio image. For time series data, ratio images were corrected for photobleaching as described previously^38,40^. When creating pseudo-color images, the lowest and highest 5% of ratio values were not used in determining the scale or the max and min colors, to eliminate spurious pixels. The images were normalized so that the lowest value was 1.

### Cross-correlation analysis pipeline

Edge-aligned GTPase activity and edge velocity measurements were obtained using a previously described windowing and edge-tracking workflow (https://github.com/DanuserLab/u-integrate; https://github.com/DanuserLab/u-register)^20,41^. Briefly, the cell boundary was segmented for each frame and used to propagate edge-aligned sampling windows over time, generating five layers orthogonal to the edge. Unless otherwise noted, analysis used 20-pixel padding, light mask erosion (2 px), and five layers (PerpSize = 4 px; 0.216 µm/px). The number of windows along the boundary was selected for each movie from the median cell perimeter to achieve an approximately constant arc-length spacing. Photobleach corrected GTPase ratio images were sampled within each window to generate layer-resolved Rac1 activity maps. Edge velocity was computed from boundary displacement and sampled in the same window coordinates to generate matched velocity maps. To focus coupling analyses on protrusive regions, we classified windows as active or quiescent using the layer-1 (edge-proximal) velocity time series. For each window, we applied a Ljung–Box test on the velocity trace (maximum lags set to 50% of the series length, capped at 20; α = 0.10). Per-window p-values were smoothed along the boundary using a short moving mean, and windows with smoothed p-values below α were labeled active. Cross-correlation analyses were performed using active windows only.

### Cluster permutation analysis

To compare WT and perturbations across the lag, we used a cluster-based permutation test^42^. Cross-correlation values were Fisher z–transformed and a pointwise Welch’s t-test was applied at each lag. Adjacent lags that exceeded the threshold (p < 0.05) were grouped into clusters (minimum cluster size = 3 time points), and cluster strength was summarized by the sum of the absolute t-statistics within each cluster (“cluster mass”). To control family-wise error across the tested lag window, permutation testing was performed by randomly shuffling cell labels between WT and comparison group 5000 times. For each permutation, Welch’s t-test were recalculated at each lag, clusters were identified using the same threshold and minimum cluster size, and the maximum cluster mass observed in that permutation was recorded to generate the null distribution. Clusters were considered significant at a family-wise error rate of p < 0.05. Testing was restricted to the positive-lag range (0-2 min).

### Protrusion-onset–aligned Rac1 activity

To quantify Rac1 activity dynamics after protrusion onset, we performed event-triggered averaging using edge-velocity time series and layer-resolved Rac1 biosensor signals sampled in edge-aligned windows^43,44^. For each cell, we used only active windows (as defined by Ljung-Box test)^39^. For each window, edge velocity was smoothed using cubic smoothing splines (csaps, smoothing parameter = 0.9). Protrusion phases were defined as contiguous time intervals where smoothed velocity was positive and retained only if their duration exceeded a minimum phase length (≥25 s). Protrusion onset was defined as the first frame of each qualifying protrusion run. Rac1 activity was z-scored within each window to reduce variability across windows and cells. For each protrusion onset event, we extracted time-aligned segments of z-scored Rac1 activity spanning a symmetric window around onset (±50 s). Event-triggered Rac1 activity traces were then averaged across events and subsequently across cells for each layer to obtain the population mean response, with uncertainty reported as 2×SEM across events.

### Cell motion summary metrics

Velocity time series were analyzed in active windows defined by the Ljung–Box test. For each active window, edge velocity was denoised using a 3-frame running median filter. Protrusions were identified from the filtered velocity using a hysteresis threshold to suppress single-frame sign flips: protrusions were initiated when velocity exceeded 50 nm s⁻¹ and terminated when it fell below 25 nm s⁻¹. Events shorter than 10 s were excluded. Protrusions were defined as contiguous suprathreshold runs of frames. Protrusion fraction was calculated for each window as the fraction of frames classified as protrusive, and was then averaged across active windows within each cell. Retraction fraction was calculated analogously, with retraction initiated when velocity decreased below −50 nm s⁻¹ and terminated when it rose above −25 nm s⁻¹. To quantify directed migration behavior, centroid of the cell mask was tracked over time to measure both total displacement and net displacement. Directed migration was defined as the ratio of net displacement to total displacement.

## Supporting information

Rac1 activity reported by AlloRac1 in a randomly migrating MEF cell (left). Colored lines show the cell edge at successive times (right).

Cdc42 activity reported by AlloCdc42 in a randomly migrating MEF cell (left). Colored lines show the cell edge at successive times (right).

## Acknowledgements

KH thanks the NIGMS for financial support (grant R35-GM122596). MC thanks the NSF GRFP and MiBio T32 Training grants for financial support. We thank the Janelia Research Campus for providing the dyes used in this study (https://janeliamaterials.azurewebsites.net). NVS-Pak1 was a kind gift from Dr. David Wolff (Augusta University)

## Author contributions

MC initiated the project and performed the bulk of the experiments and analysis. RR assisted with computational analysis, including tests for potential roles of Rac1 feedback loops. LH carried out FACS sorting, and helped with generation of stable cell lines and Western blotting. YY helped with optimization of the Cdc42 chromophore side chain interactions. DL assisted with cloning and molecular biology. GD and KH directed the project and provided intellectual input. MC and KH wrote the paper, with input from all authors.

## Data and code availability statement

The datasets generated or analyzed during and/or analyzed during the study are available from the corresponding author on reasonable request.

## Extended data figures and table

**Extended Data Figure 1.**
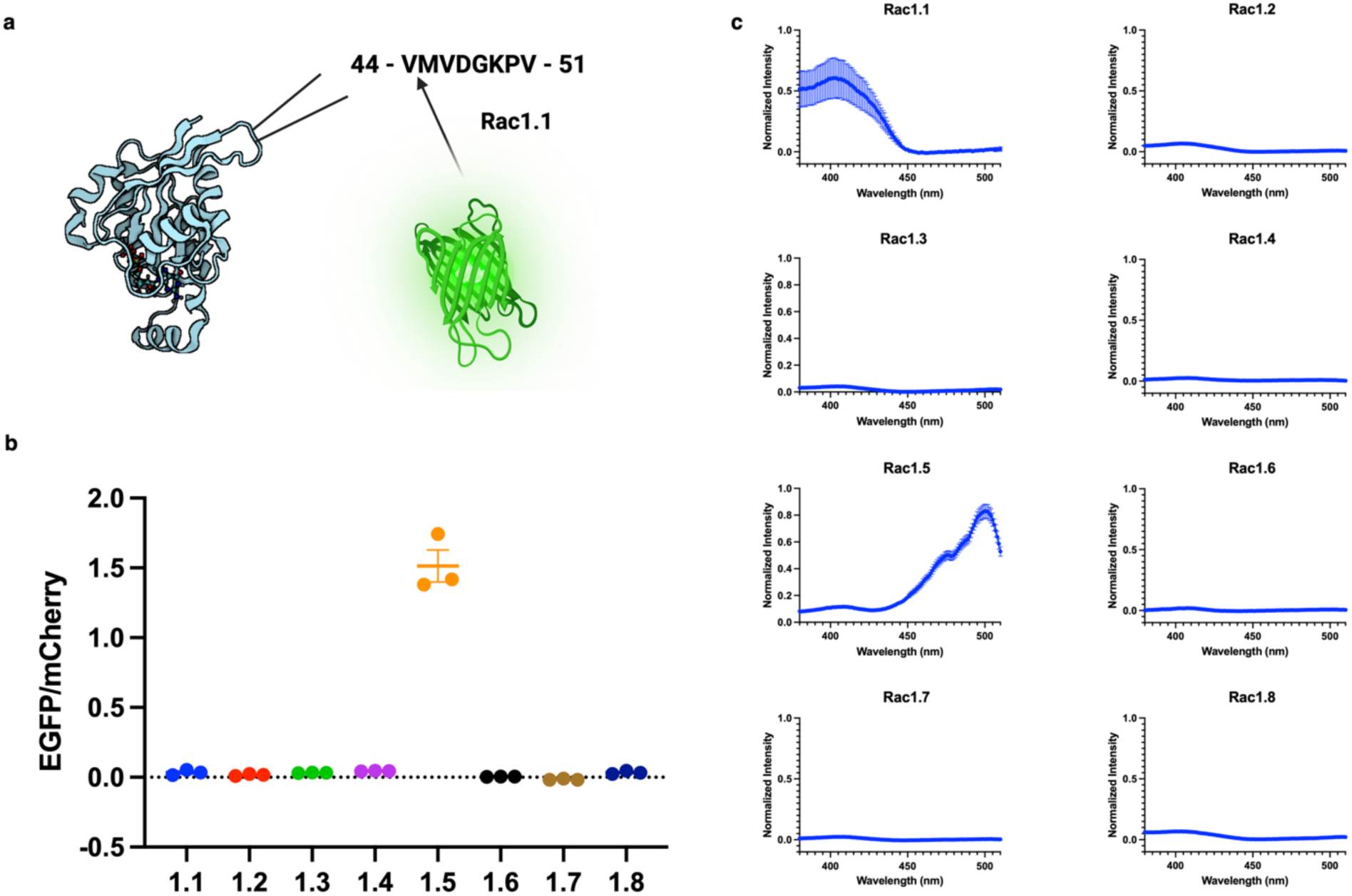
Screening cpGFP insertion positions within the Rac1 allosteric loop. **a,** Circularly permuted GFP (cpGFP) was inserted after each residue of the Rac1 allosteric loop (Rac1 left). Constructs are named by insertion position (for example, Rac1.1 indicates cpGFP inserted after the first residue of the loop, Rac1.2 after the second residue, etc.). **b,** Fluorescence output of each insertion variant expressed in HEK293T cells. (Cells were excited at 480 nm. Graph shows the ratio of peak GFP emission to peak mCherry emission. n = 3 wells per condition; error bars, s.d). **c,** Excitation spectra for all insertion variants measured with emission fixed at 520 nm. Data represent n = 3 wells per condition; error bars, s.d. For **b** and **c**, each construct was fused to mCherry to normalize for expression level. Spectra of untransfected cells were used for background subtraction.

**Extended Data Figure 2.**
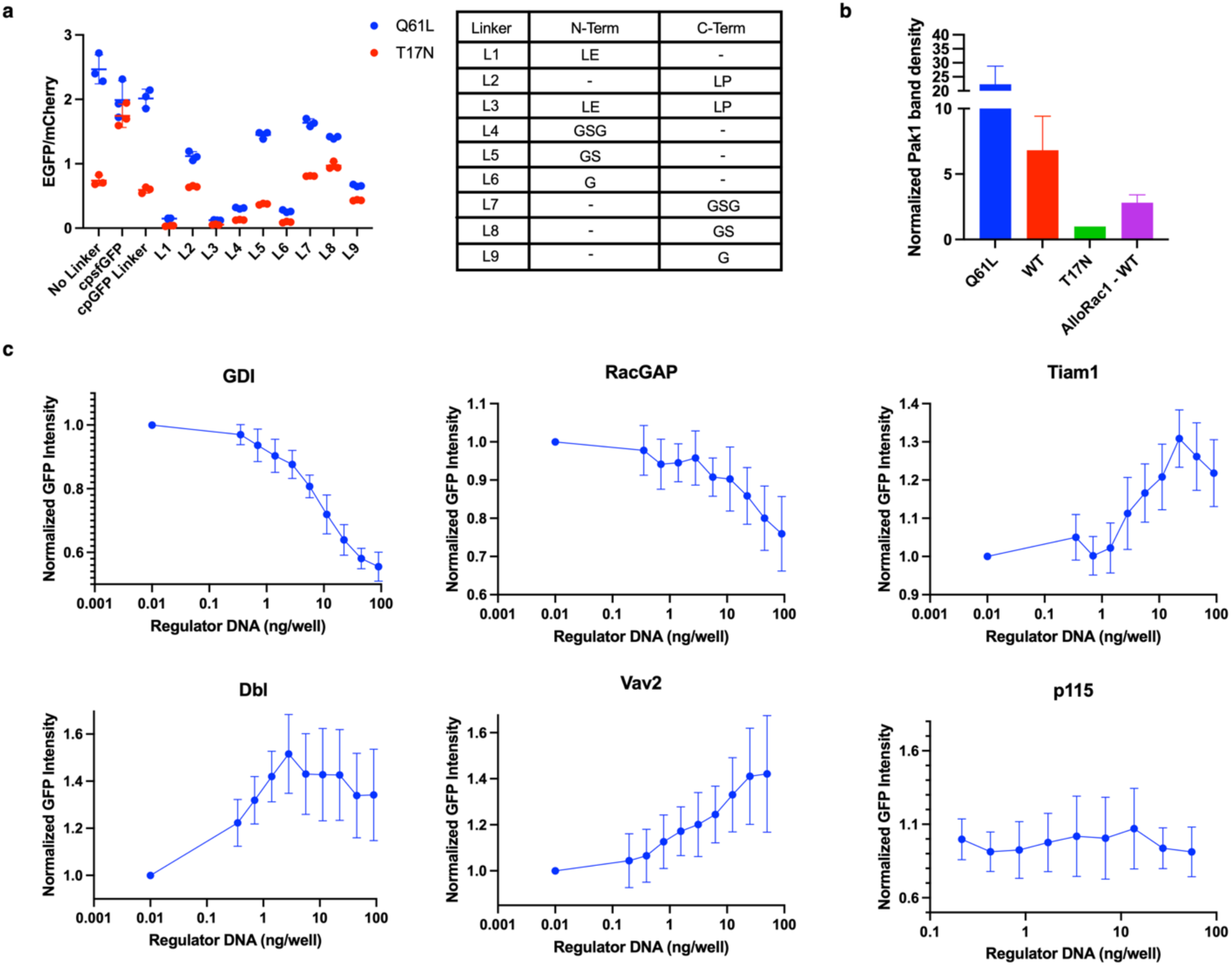
Optimization and validation of AlloRac1. **a,** Fluorescence of linker and fluorophore variants. Constructs were fused to mCherry and expressed in HEK293T cells. Active (Q61L) and inactive (T17N) mutants were compared to gauge fluorescence response to conformational state. Cells were excited at 480 nm, GFP fluorescence was normalized for expression using mCherry, and the output is reported as the ratio of peak GFP emission to peak mCherry emission, after background subtraction. The background was determined using untransfected cells. The table indicates the linker combinations tested (positions labeled relative to cpGFP N- and C-termini). cpsfGFP denotes circularly permuted superfolder GFP (Addgene #102550). cpGFP linker indicates insertion of an additional glycine in the linker joining the original GFP N- and C-termini. Data represent n = 3 wells per condition; error bars, s.d. **b,** Quantifying co-immunoprecipitation of endogenous Pak1 with Rac1 constructs in HEK293T cells. Values represent Pak1 band density normalized to Rac1 band density in the eluted fraction. Data represent N = 3 independent experiments; error bars, s.d. **c,** Titration of upstream regulators in HEK293T cells and corresponding changes in AlloRac1 activity. GFP fluorescence was normalized to HaloTag–JF646 to account for expression level, and responses were then normalized to conditions with no regulator (first titration point). At the highest concentrations of some regulators, we observed effects on cell morphology . These wells were not included in the analysis. N = 3 independent experiments with n = 12 technical replicates per condition; for p115, N = 2 and n = 5. Error bars, s.d.

**Extended Data Figure 3.**
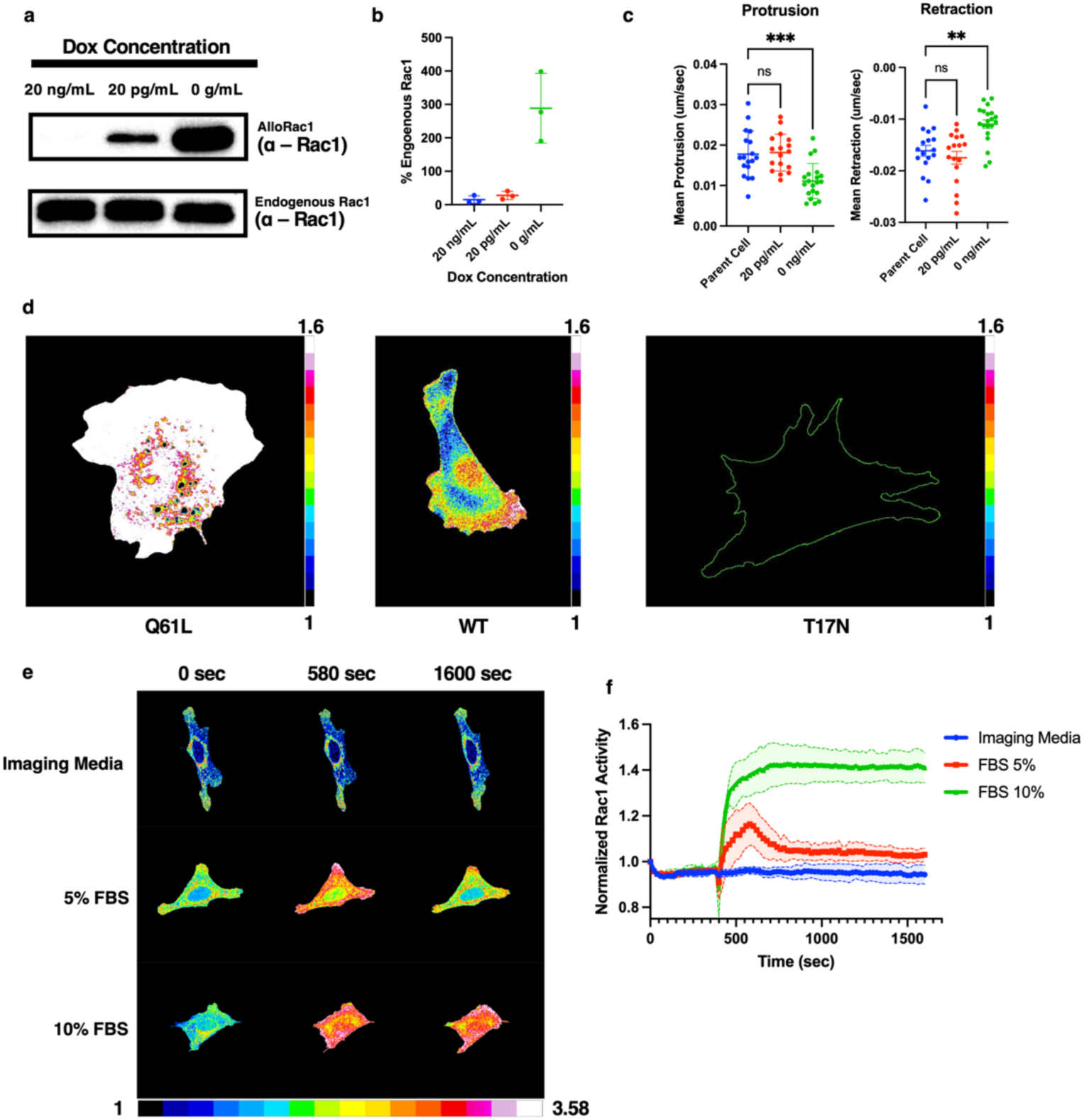
Characterization of a stable MEF AlloRac1 cell line. **a,** Representative western blot showing AlloRac1 expression relative to endogenous Rac1 across doxycycline concentrations. Rac1 was detected using an anti-Rac1 antibody (Cell Signaling Technology #2465). **b,** Quantification of AlloRac1 expression relative to endogenous Rac1 in the stable MEF cell line. Values represent AlloRac1 band density normalized to endogenous Rac1 band density and reported as percent of endogenous Rac1. N = 3 independent experiments; error bars, s.d. **c,** Protrusion dynamics in the stable AlloRac1 MEF line induced with low (20 pg/mL) or high (20 ng/mL) doxycycline, compared to the parental (WT) line. A membrane marker was expressed in parental cells to enable matched imaging conditions. Each dot represents one cell. N = 2 independent experiments with n = 18 (WT), n = 17 (low), and n = 20 (high); error bars, s.d. Ordinary one-way ANOVA; ns, P > 0.05; **P ≤ 0.01; ***P ≤ 0.001 **d,** Representative Rac1 activity imaged in MEFs expressing constitutively active (Q61L), wild-type, or inactivated (T17N) AlloRac1, acquired under identical imaging conditions. Images are contrast-matched to the wild-type condition. **e,** Time-lapse montage of Rac1 activity before and after serum stimulation at the indicated serum concentrations. Cells were serum-starved (0.1% FBS, 24 h) and imaged at 20 second intervals, for 20 frames before and 60 frames after stimulation. Images were processed and corrected for photobleaching. Images are contrast-matched. **f,** Quantification of mean Rac1 activity over time. Curves show the population mean, with shaded regions indicating s.d.; traces are normalized to the first time point. n = 6 cells (imaging media), n = 3 cells (5% FBS), and n = 3 cells (10% FBS).

**Extended Data Figure 4.**
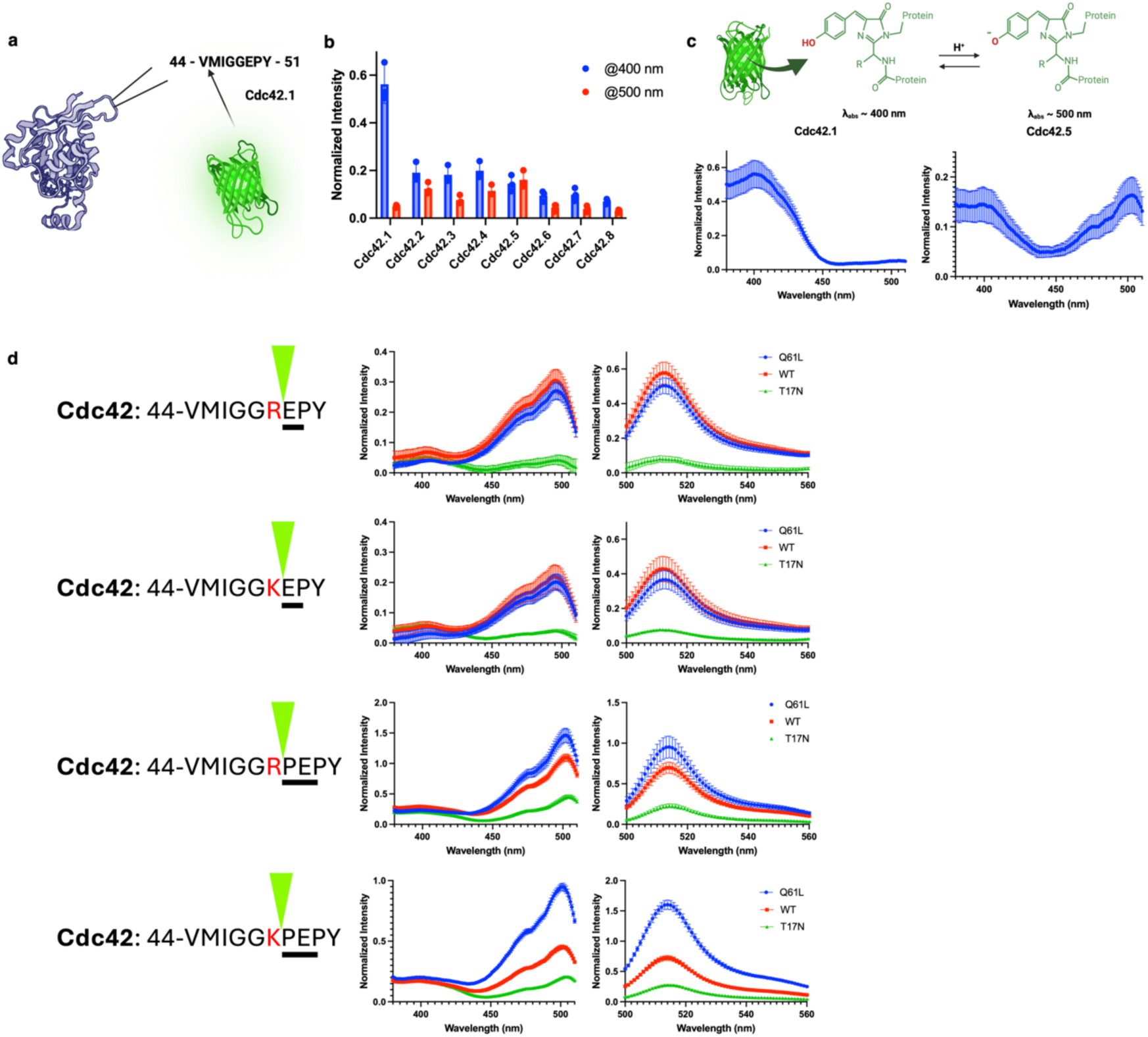
Optimization of AlloCdc42. **a,** cpGFP was inserted after each residue in the Cdc42 allosteric loop (Cdc42 left). Constructs are named by insertion position as in Supplementary Fig.1. **b,** Fluorescence output of each insertion variant expressed in HEK293T cells. Constructs were fused to mScarlet to control for expression level. Excitation spectra were acquired with emission fixed at 520 nm, background-subtracted using untransfected cells, and normalized to the peak mScarlet emission. Normalized values at 400 nm and 500 nm (from the two excitation peaks) are reported. n = 3 wells per condition; error bars, s.d. **c,** Schematic of the GFP fluorophore illustrating two states associated with the two excitation peaks, and excitation spectra for two insertion variants; One is dominated by a single fluorophore state, and the other shows prominent contributions from both states. Lines show the mean of n = 3 wells per condition; error bars, s.d. **d,** Optimizing the local environment around the cpGFP fluorophore. The cpGFP insertion position is indicated (green arrow), and the neighboring charged residue is highlighted (red). The corresponding Cdc42 allosteric loop sequence is shown, along with excitation and emission spectra of Cdc42 constructs in constitutively active (Q61L), wild-type, and inactivated (T17N) states, expressed in HEK293T cells. GFP fluorescence was normalized to either mScarlet or HaloTag-JF646 to account for expression level. Underlines indicate inserted residues. Lines show the mean of n = 3 wells per condition; error bars, s.d.

**Extended Data Figure 5.**
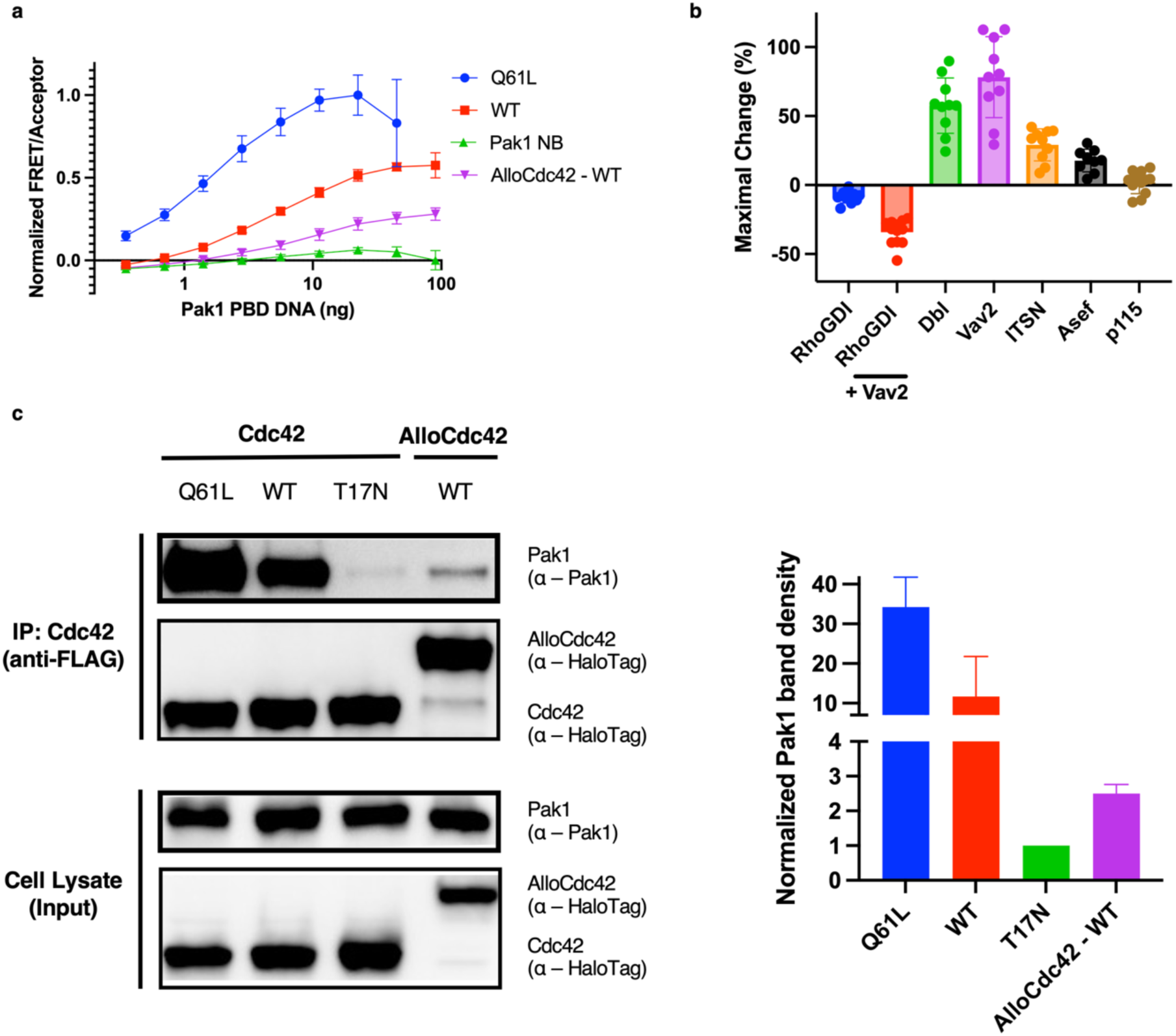
Validation of AlloCdc42. **a,** FRET-based binding assay for AlloCdc42, used in (Fig. 1c). Data represent n = 6 technical replicates per condition; error bars, s.d. **b,** Titration of upstream regulators in HEK293T cells and corresponding changes in AlloCdc42 activity. For the RhoGDI+Vav2 condition, cells were co-expressed with Vav2 to activate Cdc42 before RhoGDI titration. Responses are reported as maximal percent change relative to baseline (no regulator). N = 3 independent experiments with n = 10 (RhoGDI), n = 12 (RhoGDI + Vav2), n = 10 (Dbl), n = 10 (Vav2), n = 10 (ITSN), n = 9 (Asef), and n = 12 (p115) technical replicates per condition; error bars, s.d. **c,** Representative co-immunoprecipitation showing association of Cdc42 constructs with endogenous Pak1 in HEK293T cells (left). Co-immunoprecipitation of endogenous Pak1 with Cdc42 constructs in HEK293T cells. Values represent Pak1 band density normalized to Cdc42 band density in the eluted fraction. N = 2 independent experiments; error bars, s.d. (right).

**Extended Data Figure 6.**
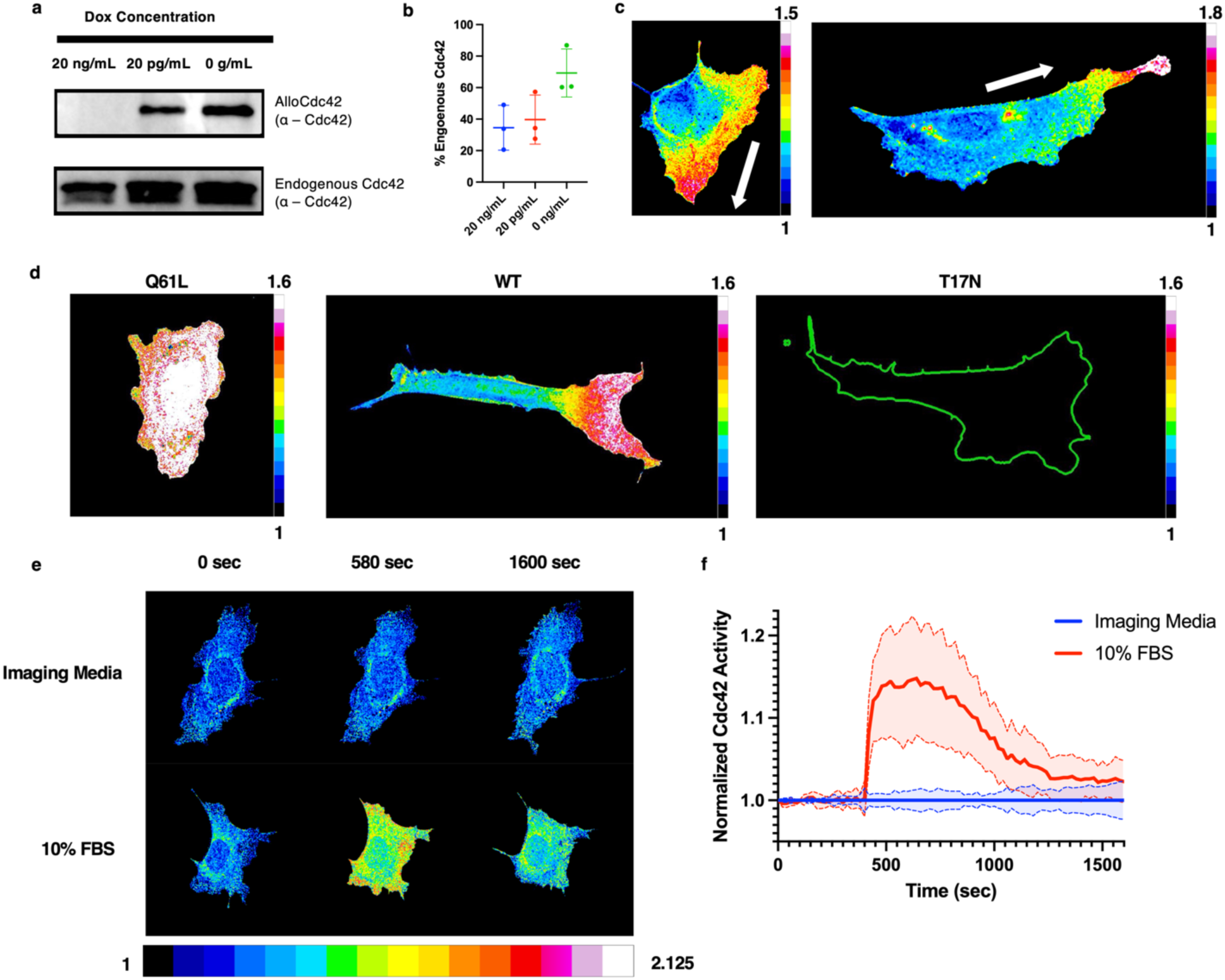
Characterization of a stable MEF AlloCdc42 cell line. **a,** Representative western blot showing AlloCdc42 expression relative to endogenous Cdc42 across doxycycline concentrations. Cells were selected using puromycin, then sorted for AlloCdc42 expression using FACS. Cdc42 was detected using an anti-Cdc42 antibody (Cell Signaling Technology #2462). **b,** Quantification of AlloCdc42 expression relative to endogenous Cdc42 in the stable MEF cell line. Values represent AlloCdc42 band density normalized to endogenous Cdc42 band density and reported as percent of endogenous Cdc42. Data represent N = 3 independent experiments; error bars, s.d. **c,** Representative MEFs, showing Cdc42 activity localized at the leading edge in cells undergoing directed motility. White arrows indicate direction of motion. **d,** Comparison of Cdc42 activity in MEFs expressing constitutively active (Q61L), wild-type, or dominant-negative (T17N) AlloCdc42, imaged under identical conditions. Image contrast and scaling was matched to the wild-type condition. **e,** Cdc42 activity before and after serum stimulation at the indicated serum concentrations. Cells were serum-starved (0.1% FBS, 24 h) and imaged for 20 frames before and 60 frames after stimulation, at 20 second intervals. Images were photobleach-corrected and contrast-matched. **f,** Quantification of mean Cdc42 activity over time. Curves show the population mean, with shaded regions indicating s.d.; traces are normalized to the first time point. n = 3 cells (imaging media), n = 3 cells (10% FBS).

**Extended Data Figure 7.**
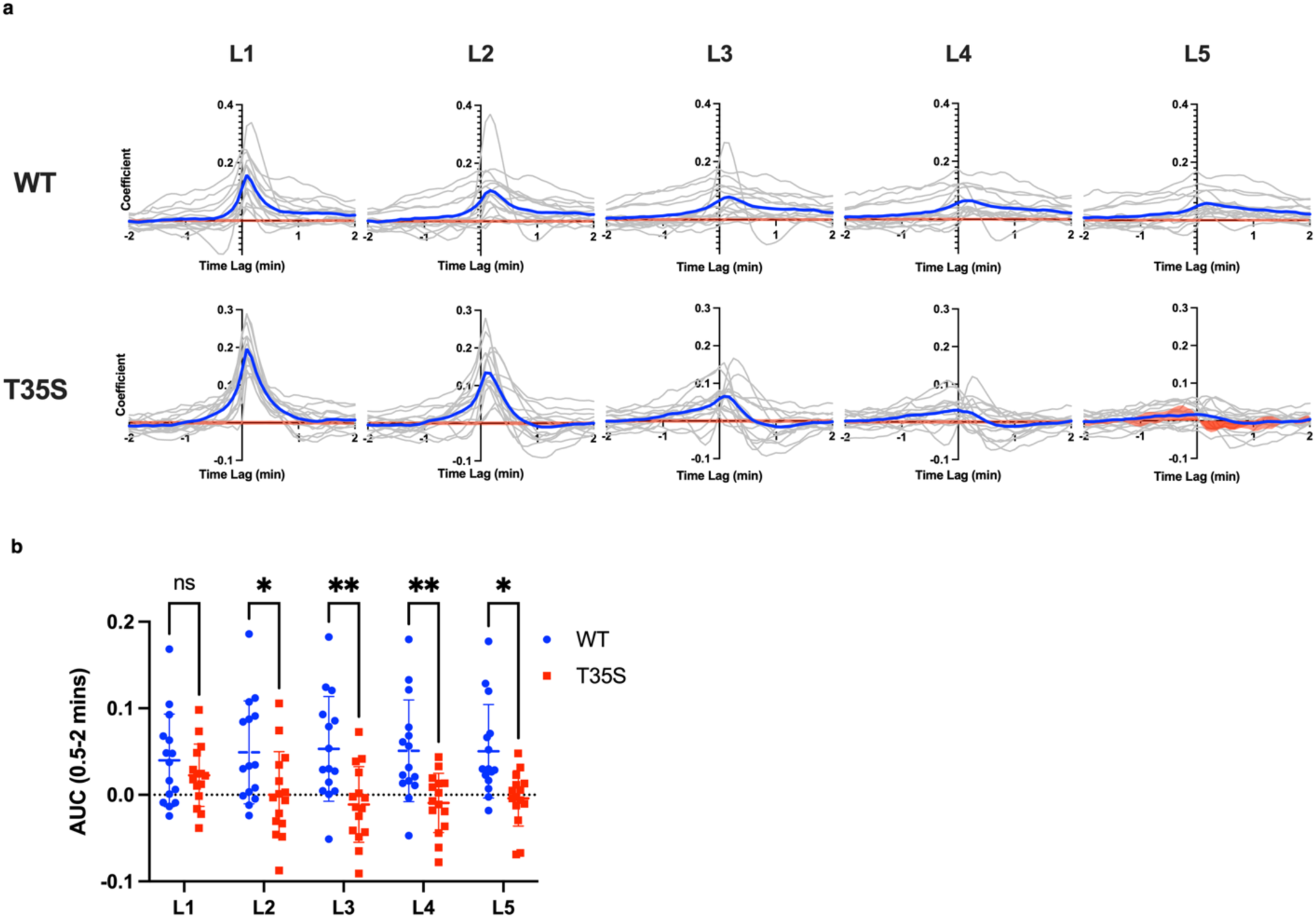
Layer-dependent Rac1–edge cross-correlation analysis. **a,** Cross-correlation curves for individual cells (grey). Blue: average cross-correlation curve for the population in each layer (from N = 3 independent experiments with n = 15 cells per condition). Red shaded areas indicate 95% permutation-based confidence level from time-shuffled data (1000 times). **b,** Area under the curve (AUC) computed as the trapezoidal integral of the cross-correlation curve over the positive-lag (0.5-2min). Same population as (**a**). Ordinary two-way ANOVA, Šídák’s multiple comparisons test ; ns, P > 0.05; *P ≤ 0.05; **P ≤ 0.01

**Extended Data Figure 8.**
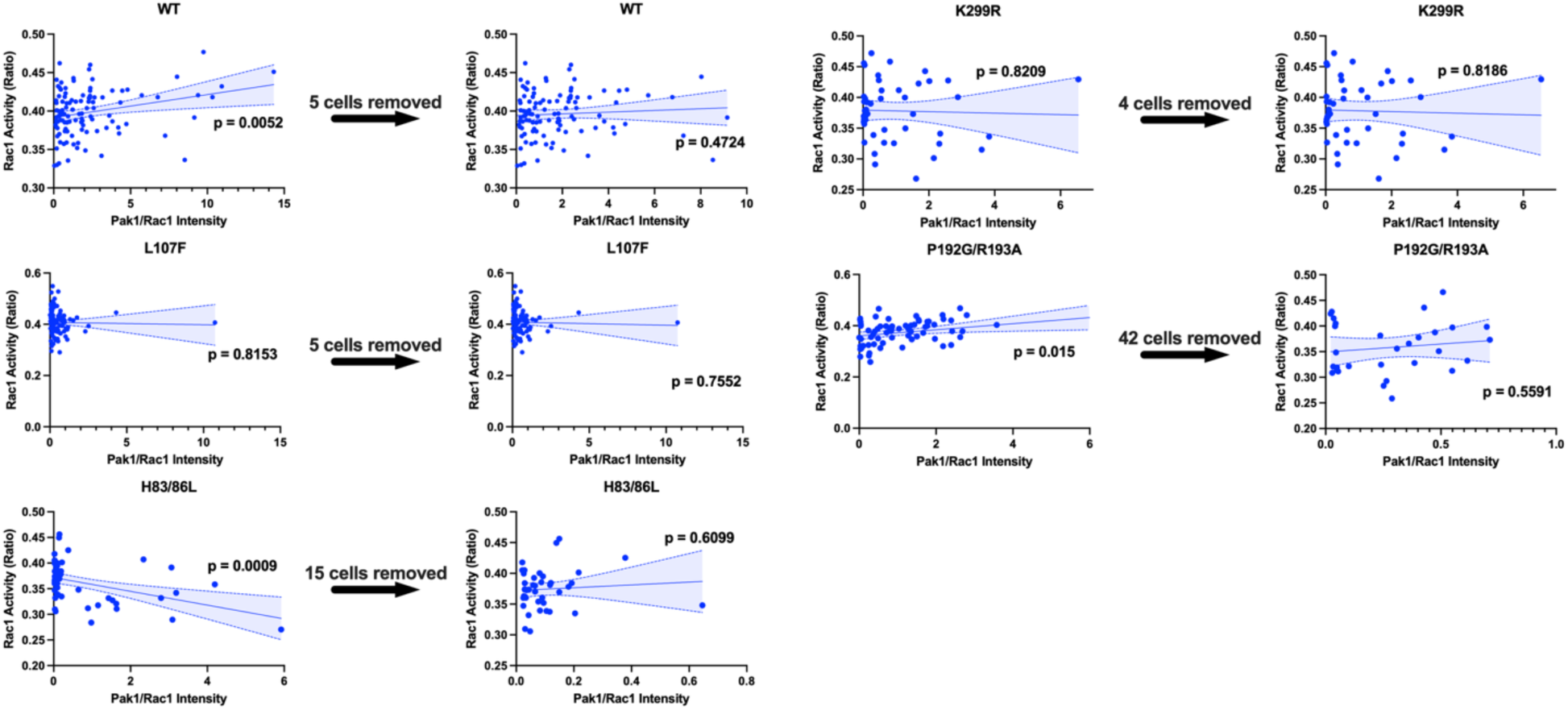
Relationship between Pak1 expression, Rac1 expression and Rac1 activity. Pak1/Rac1 expression, determined from mean intensity ratios, is plotted against Rac1 activity (mean biosensor ratio per cell) before and after applying selection criteria. Cells were first filtered for minimum Pak1 expression (including only cells with mean intensity >100 a.u.). Cells with the highest Pak1/Rac1 ratios were then sequentially excluded until the relationship between the Pak1/Rac1 ratio and Rac1 activity was no longer significant (p > 0.4). Points represent individual cells; lines indicate linear regression with 95% confidence bands (p-value for nonzero slope).

**Extended Data Figure 9.**
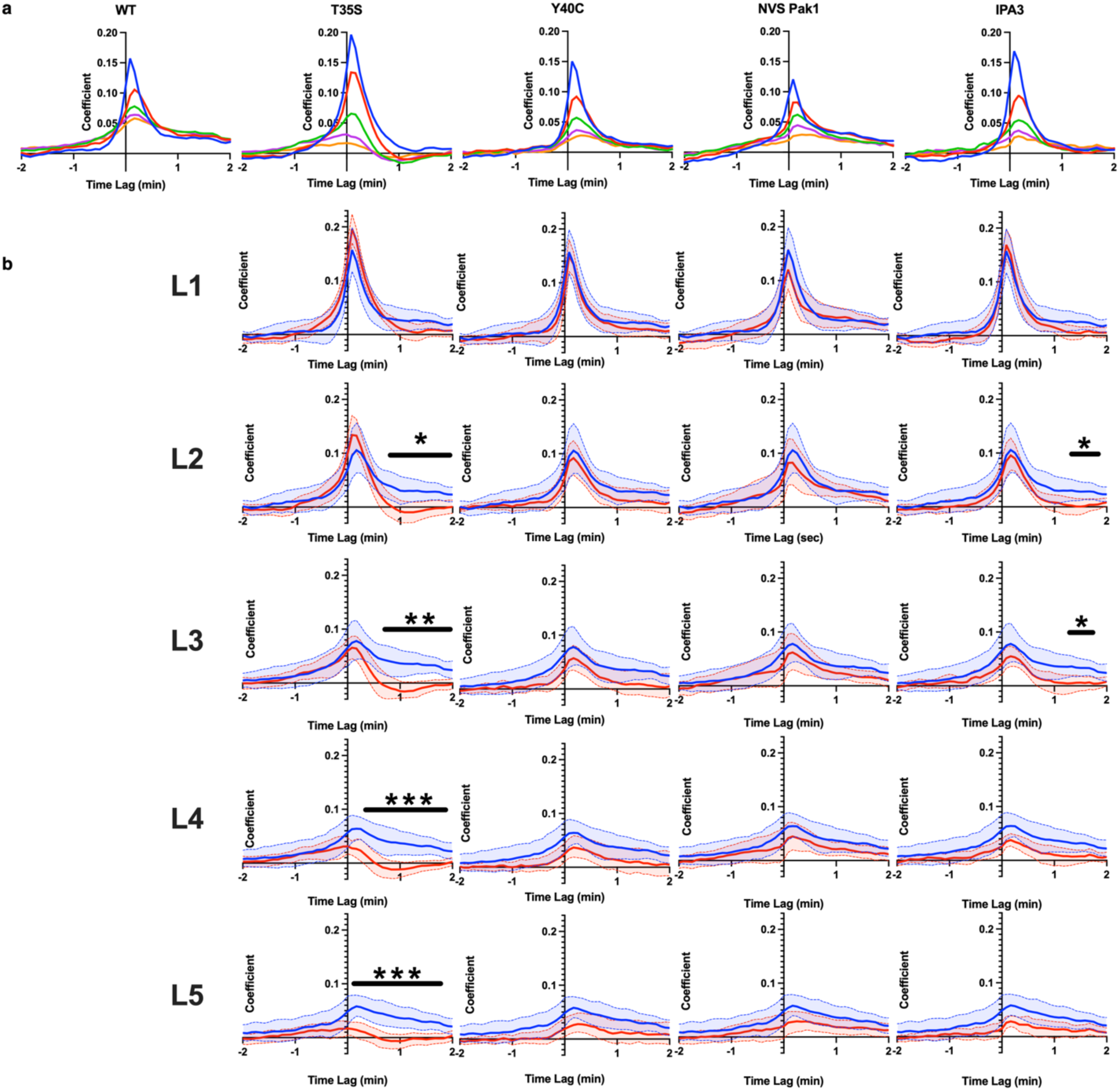
Effects of Rac1 mutations and inhibitors on the correlation between Rac1 activity and edge motion. **a,** Cross-correlation between Rac1 activity and edge velocity for the downstream effector mutants and Pak1 inhibitor conditions used in Fig. 4b. Curves from all edge layers are superimposed. WT and T35S are reproduced from Fig. 3e to facilitate comparison with the additional conditions shown here. **b,** Cluster-based permutation analysis of cross-correlation curves for each condition (red) separated by edge layer. For each condition, the results of mutations and inhibitors are shown in red and the WT curves in blue. WT and T35S are duplicated from Fig. 3f. to facilitate comparison across all conditions. Black bars denote suprathreshold clusters; significance is indicated as (*P ≤ 0.05; **P ≤ 0.01; ***P ≤ 0.001).

**Extended Data Table 1:** AlloGTPase Sensor Brightness (QY x ε, see methods)

| Sensor | Brightness |
| --- | --- |
| AlloRac1 | 31,9500 ± 1,560 |
| AlloCdc42 | 11,400 ± 480 |

## Supplementary Information

Supplementary Movie 1

Supplementary Video 2

**Supplementary Table 1:** Important plasmids constructed and used for this study.

| Plasmids | Insert 1 | Insert 2 | Insert 3 | Vector | Source |
| --- | --- | --- | --- | --- | --- |
| HaloTag cpEGFP Rac1.5 Q61L | HaloTag | Rac1 | cpEGFP | pEGFP-C1 | This study |
| HaloTag cpEGFP Rac1.5 WT | HaloTag | Rac1 | cpEGFP | pEGFP-C1 | This study |
| HaloTag cpEGFP Rac1.5 T17N | HaloTag | Rac1 | cpEGFP | pEGFP-C1 | This study |
| FLAG-HaloTag cpEGFP Rac1.5 Q61L | HaloTag | Rac1 | cpEGFP | pEGFP-C1 | This study |
| FLAG-HaloTag cpEGFP Rac1.5 WT | HaloTag | Rac1 | cpEGFP | pEGFP-C1 | This study |
| FLAG-HaloTag cpEGFP Rac1.5 T17N | HaloTag | Rac1 | cpEGFP | pEGFP-C1 | This study |
| HaloTag cpEGFP Cdc42.5 Q61L – KP insertion | HaloTag | Cdc42 | cpEGFP | pEGFP-C1 | This study |
| HaloTag cpEGFP Cdc42.5 WT – KP insertion | HaloTag | Cdc42 | cpEGFP | pEGFP-C1 | This study |
| HaloTag cpEGFP Cdc42.5 T17N – KP insertion | HaloTag | Cdc42 | cpEGFP | pEGFP-C1 | This study |
| FLAG-HaloTag cpEGFP Cdc42.5 Q61L – KP insertion | HaloTag | Cdc42 | cpEGFP | pEGFP-C1 | This study |
| FLAG-HaloTag cpEGFP Cdc42.5 WT – KP insertion | HaloTag | Cdc42 | cpEGFP | pEGFP-C1 | This study |
| FLAG-HaloTag cpEGFP Cdc42.5 T17N – KP insertion | HaloTag | Cdc42 | cpEGFP | pEGFP-C1 | This study |
| FLAG-HaloTag cpEGFP Rac1.5 Q61L | HaloTag | Rac1 | cpEGFP | Piggybac (Puro) | This study |
| FLAG-HaloTag cpEGFP Rac1.5 WT | HaloTag | Rac1 | cpEGFP | Piggybac (Puro) | This study |
| FLAG-HaloTag cpEGFP Rac1.5 T17N | HaloTag | Rac1 | cpEGFP | Piggybac (Puro) | This study |
| FLAG-HaloTag cpEGFP Rac1.5 T35S | HaloTag | Rac1 | cpEGFP | Piggybac (Puro) | This study |
| FLAG-HaloTag cpEGFP Rac1.5 Y40C | HaloTag | Rac1 | cpEGFP | Piggybac (Puro) | This study |
| HaloTag cpEGFP Cdc42.5 Q61L – KP insertion | HaloTag | Cdc42 | cpEGFP | Piggybac (Puro) | This study |
| HaloTag cpEGFP Cdc42.5 WT – KP insertion | HaloTag | Cdc42 | cpEGFP | Piggybac (Puro) | This study |
| HaloTag cpEGFP Cdc42.5 T17N – KP insertion | HaloTag | Cdc42 | cpEGFP | Piggybac (Puro) | This study |

**Supplementary Table 2:**
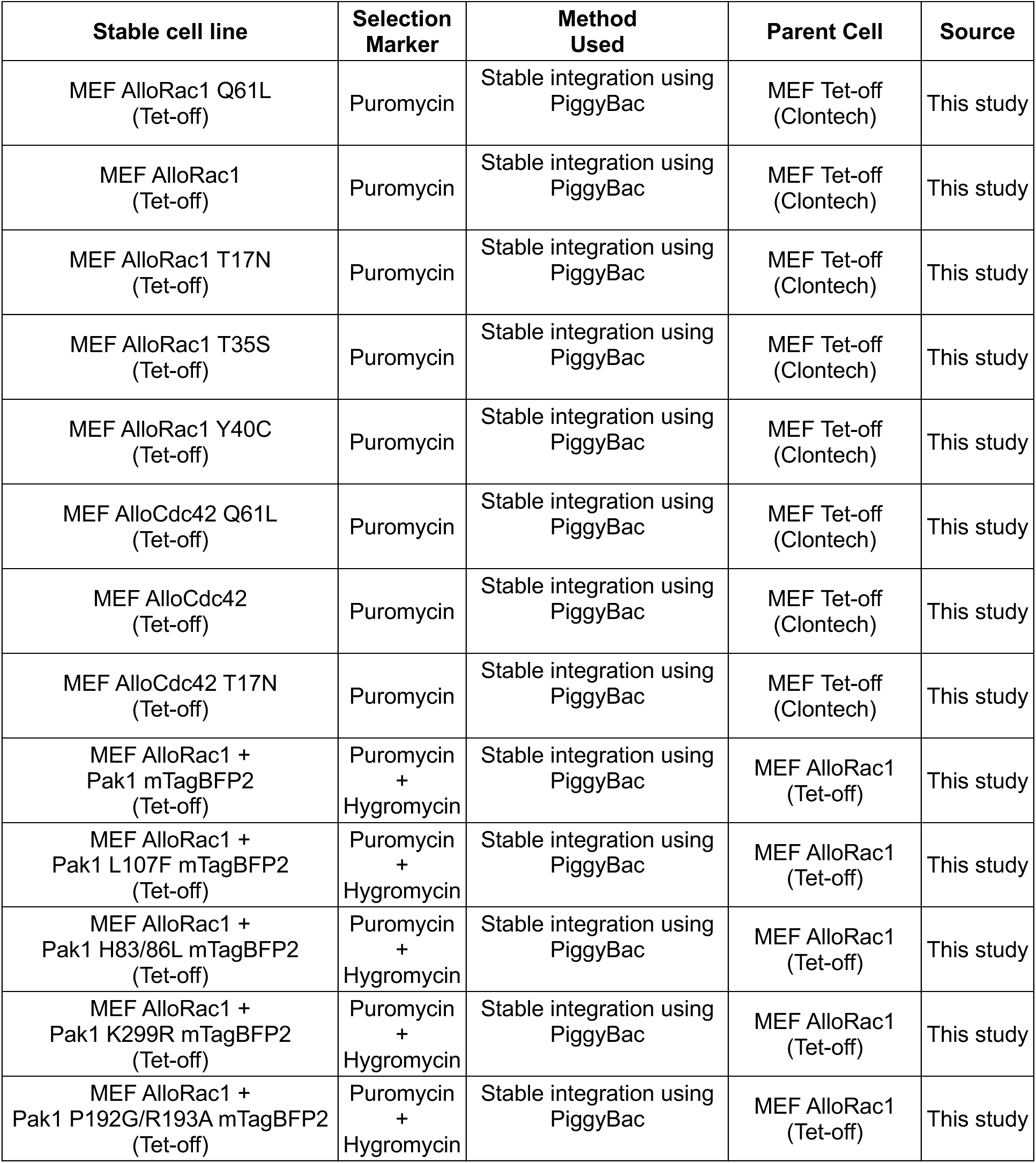
Stable cell lines used for this study.

| **Stable cell line** | **Selection Marker** | **Method Used** | **Parent Cell** | **Source** |
| --- | --- | --- | --- | --- |
| MEF AlloRac1 Q61L (Tet-off) | Puromycin | Stable integration using PiggyBac | MEF Tet-off (Clontech) | This study |
| MEF AlloRac1 (Tet-off) | Puromycin | Stable integration using PiggyBac | MEF Tet-off (Clontech) | This study |
| MEF AlloRac1 T17N (Tet-off) | Puromycin | Stable integration using PiggyBac | MEF Tet-off (Clontech) | This study |
| MEF AlloRac1 T35S (Tet-off) | Puromycin | Stable integration using PiggyBac | MEF Tet-off (Clontech) | This study |
| MEF AlloRac1 Y40C (Tet-off) | Puromycin | Stable integration using PiggyBac | MEF Tet-off (Clontech) | This study |
| MEF AlloCdc42 Q61L (Tet-off) | Puromycin | Stable integration using PiggyBac | MEF Tet-off (Clontech) | This study |
| MEF AlloCdc42 (Tet-off) | Puromycin | Stable integration using PiggyBac | MEF Tet-off (Clontech) | This study |
| MEF AlloCdc42 T17N (Tet-off) | Puromycin | Stable integration using PiggyBac | MEF Tet-off (Clontech) | This study |
| MEF AlloRac1 + Pak1 mTagBFP2 (Tet-off) | Puromycin + Hygromycin | Stable integration using PiggyBac | MEF AlloRac1 (Tet-off) | This study |
| MEF AlloRac1 + Pak1 L107F mTagBFP2 (Tet-off) | Puromycin + Hygromycin | Stable integration using PiggyBac | MEF AlloRac1 (Tet-off) | This study |
| MEF AlloRac1 + Pak1 H83/86L mTagBFP2 (Tet-off) | Puromycin + Hygromycin | Stable integration using PiggyBac | MEF AlloRac1 (Tet-off) | This study |
| MEF AlloRac1 + Pak1 K299R mTagBFP2 (Tet-off) | Puromycin + Hygromycin | Stable integration using PiggyBac | MEF AlloRac1 (Tet-off) | This study |
| MEF AlloRac1 + Pak1 P192G/R193A mTagBFP2 (Tet-off) | Puromycin + Hygromycin | Stable integration using PiggyBac | MEF AlloRac1 (Tet-off) | This study |

